# Virtual connectomic datasets in Alzheimer’s Disease and aging using whole-brain network dynamics modelling

**DOI:** 10.1101/2020.01.18.911248

**Authors:** Lucas Arbabyazd, Kelly Shen, Zheng Wang, Martin Hofmann-Apitius, Petra Ritter, The Alzheimer’s Disease Neuroimaging Initiative, Anthony R. McIntosh, Demian Battaglia, Viktor Jirsa

**Affiliations:** Université Aix-Marseille, INSERM UMR 1106, Institut de Neurosciences des Systèmes, F-13005 Marseille, France; Rotman Research Institute, Baycrest Centre, Toronto, Ontario, M6A 2E1, Canada; Fraunhofer Institute for Algorithms and Scientific Computing, 53754 Sankt Augustin, Germany; Brain Simulation Section, Department of Neurology, Charité University Medicine Berlin & Berlin Institute of Health, Germany; Bernstein Center for Computational Neuroscience Berlin, Germany; Einstein Center for Neuroscience Berlin, Charitéplatz 1, 10117 Berlin; Einstein Center Digital Future, Wilhelmstraße 67, 10117 Berlin; University of Strasbourg Institute for Advanced Studies (USIAS), 67000 Strasbourg, France

## Abstract

Large neuroimaging datasets, including information about structural (SC) and functional connectivity (FC), play an increasingly important role in clinical research, where they guide the design of algorithms for automated stratification, diagnosis or prediction. A major obstacle is, however, the problem of missing features (e.g., lack of concurrent DTI SC and resting-state fMRI FC measurements for many of the subjects).

We propose here to address the missing connectivity features problem by introducing strategies based on computational whole-brain network modeling. Using two datasets, the ADNI dataset and a healthy aging dataset, for proof-of-concept, we demonstrate the feasibility of virtual data completion (i.e., inferring “virtual FC” from empirical SC or “virtual SC” from empirical FC), by using self-consistent simulations of linear and nonlinear brain network models. Furthermore, by performing machine learning classification (to separate age classes or control from patient subjects) we show that algorithms trained on virtual connectomes achieve discrimination performance comparable to when trained on actual empirical data; similarly, algorithms trained on virtual connectomes can be used to successfully classify novel empirical connectomes. Completion algorithms can be combined and reiterated to generate realistic surrogate connectivity matrices in arbitrarily large number, opening the way to the generation of virtual connectomic datasets with network connectivity information comparable to the one of the original data.

**Significance statement:** Personalized information on anatomical connectivity (“structural connectivity”, SC) or coordinated resting state activation patterns (“functional connectivity’, FC) is a source of powerful neuromarkers to detect and track the development of neurodegenerative diseases. However, there are often “gaps” in the available information, with only SC (or FC) being known but not FC (or SC). Exploiting whole-brain modelling, we show that gap in databases can be filled by inferring the other connectome through computational simulations. The generated virtual connectomic data carry information analogous to the one of empirical connectomes, so that machine learning algorithms can be trained on them. This opens the way to the release in the future of cohorts of “virtual patients”, complementing traditional datasets in data-driven predictive medicine.

## Introduction

One of the greatest challenges today is to develop approaches allowing the useful exploitation of large-scale datasets in biomedical research in general (Margolis et al., 2014) and neuroscience and neuroimaging in particular (Van Horn and Toga, 2014). Progress in this direction is made possible by the increasing availability of large public datasets in the domain of connectomics (Van Essen et al., 2013; Poldrack and Gorgolewski, 2014; Horien et al., 2020). This is true, in particular, for research in Alzheimer’s disease (AD), in which, despite decades of massive investment and a daunting literature on the topic, the partial and, sometimes contradictory nature of the reported results (World Alzheimer Report 2018) still prevents a complete understanding of the factors governing the progression of the disease (Braak & Braak, 1991; Braak et al., 2006; Komarova & Thalhauser, 2011; Henstridge et al., 2019) or of the diversity of cognitive deficits observed in different subjects (Iacono et al., 2009; Mungas et al., 2010; Allen et al., 2016). In AD research, datasets that compile rich and diverse genetic, biomolecular, cognitive, and neuroimaging (structural and functional) features for a large number of patients are playing an increasingly important role (Rathore et al., 2017; Iddi et al., 2019). Example applications include: the early diagnosis and prognosis by using MRI images (Dennis & Thompson, 2014; Chiesa et al., 2017; De Vos et al., 2018); the use of machine learning for automated patient classification (Cuingnet et al., 2011; Zhang et al., 2012; Moore et al., 2019) or prediction of the conversion from early stages to fully developed AD (Rombouts et al., 2005; Moradi et al., 2015; Casanova et al., 2018), with signs of pathology difficult to distinguish from “healthy aging” effects (Doan et al., 2017); the extraction of decision networks based on the combination of semantic knowledge bases and data mining of the literature (Sanchez et al., 2011; Kodamullil et al., 2015; Iyappan et al., 2016).

Among the factors contributing to the performance of prediction and inference approaches in AD –and, more in general, other neurological or psychiatric diseases (Walter et al., 2016) or studies of aging (Cole and Franke, 2017)– are not only the large size of datasets but also the multiplicity of features jointly available for each patient. Indeed, one can take advantage not only of the complementary information that different features could bring but also capitalize on possible synergies arising from their simultaneous knowledge (Wang et al., 2015; Zimmermann et al., 2016; Iddi et al., 2019). Unfortunately, even gold standard publicly available datasets in AD, such as the datasets released by the Alzheimer’s Disease Neuroimaging Initiative (ADNI) consortium (Wyman et al., 2013; Beckett et al., 2015; Weiner et al., 2017), have severe limitations. Indeed, if they include neuroimaging features of different types –structural DTI and functional MRI– these features are simultaneously available for only a substantial minority of the subjects in the dataset (i.e., the feature coverage is not uniform over the dataset). In addition, if the number of subjects included is relatively large (hundreds of subjects), it still is too small to properly qualify as “big data”. Furthermore, the connectomic data themselves have an imperfect reliability, with a test/retest variability that can be quite large, making potentially difficult subject identifiability and, thus, personalized information extraction (Termenon et al., 2016).

Here, we will introduce a new solution aiming at relieving the problems of partially missing features and limited sample size and illustrate their validity on the two independent example datasets. Specifically, we will focus on two examples of structural and functional neuroimaging datasets, as important proofs of concept: a first one addressing AD, mediated from the previously mentioned ADNI databases (Wyman et al., 2013; Beckett et al., 2015); and a second one investigating a cohort of healthy subjects over a broad span of adult age, to analyse the effects of normal aging (Zimmermann et al., 2016; Battaglia et al., 2020). It is important to stress however that the considered issues may broadly affect any other connectomic dataset gathered for data mining intents.

To cope with missing connectomic features (and “filling the gaps” in neuroimaging datasets), we propose to build on the quickly maturating technology of mean-field whole-brain network modeling (see Deco et al., 2011 for review). Indeed, computational modeling provides a natural bridge between structural and functional connectivity, the latter emerging as the manifestation of underlying dynamical states, constrained but not entirely determined by the underlying anatomy (Ghosh et al., 2008; Kirst et al., 2016). Theoretical work has shown that average functional connectivity properties in the resting-state can be accounted for by the spontaneous collective activity of brain networks informed by empirical structural connectivity (SC) when the system is tuned to operate slightly below a critical point of instability (Deco et al., 2011, 2012). Based on this finding, simulations of a model constructed from empirical DTI connectomes and then tuned to a suitable slightly sub-critical dynamic working point are expected to provide a good rendering of resting-state functional connectivity (FC). Such whole-brain simulations are greatly facilitated by the availability of dedicated neuroinformatic platforms –such as “The Virtual Brain” (TVB; Sanz-Leon et al., 2013, 2015; Woodman et al., 2014)– and data pre-processing pipelines (Schirner et al., 2015; Proix et al., 2016), enabling brain model personalization and clinical translation (Jirsa et al., 2017; Proix et al., 2017). It thus becomes possible to complete the missing information in a dataset about BOLD fMRI FC by running a TVB simulation in the right regime, embedding the available empirical DTI SC (*SC-to-FC completion*). Analogously, algorithmic procedures based on mean-field modeling steps (“effective connectivity” approaches by Gilson et al. (2016; 2018), here used for a different purpose) can be used to address the inverse problem of inferring a reasonable ersatz of SC from resting state FC (*FC-to-SC completion*). In this study we will demonstrate the feasibility of both types of completion (SC-to-FC and FC-to-SC), applying alternative linear and nonlinear simulation pipelines to both the ADNI and the healthy ageing proof-of-concept datasets.

Beyond a single step of virtual completion, by combining completion procedures – to map, e.g., from an empirical SC (or FC) to a virtual FC (or SC) and then, yet, to a “twice virtual” SC (or FC)– we can generate for each given empirical connectome a surrogate replacement, i.e. map every empirical SC or FC to a matching *dual (bivirtual) connectome* of the same nature. We show then that pairs of empirical and bivirtual dual connectivity matrices display highly correlated network topology features, such as node-level strengths or clustering and centrality coefficients (Bullmore & Sporns, 2009). We demonstrate along the example of relevant classification tasks (stratification of mild cognitive impairment (MCI) or AD patients from control subjects on the ADNI dataset and age-class prediction on the healthy aging dataset) that close performance can be reached using machine learning algorithms trained on actual empirical connectomes or on their duals. Furthermore, empirical connectomes can be correctly categorized by classifiers trained uniquely on virtual duals.

To conclude, we provide systematic recipes for generating realistic surrogate connectomic data via data-constrained mean-field models. We show that the information that we can extract from computationally inferred connectivity matrices are only moderately degraded with respect to the one carried by the original empirical data. This opens the way to the design and sharing of veritable “virtual cohorts” data, ready for machine-learning applications in clinics, that could complement actual empirical datasets –facilitating learning through “data augmentation” (Yaeger et al., 197; Taylor & Nitschke, 2018)– or, even, potentially, fully replace them, e.g. when the sharing of real data across centers is restricted due to byzantine regulation issues (not applying to their totally synthetic but operationally-equivalent ersatz, the virtual and bivirtual duals).

## Materials and Methods

### Two datasets for proof of concept

We applied our data completion pipelines in this study to two different and independent neuroimaging datasets, from which SC and FC connectivity matrices could be extracted for at least a part of the subjects. A first dataset was obtained from the Alzheimer’s Disease Neuroimaging Initiative (ADNI) database (adni.loni.usc.edu). The ADNI was launched in 2003 as a public-private partnership, led by Principal Investigator Michael W. Weiner, MD. The primary goal of ADNI has been to test whether serial magnetic resonance imaging (MRI), positron emission tomography (PET), other biological markers, and clinical and neuropsychological assessment can be combined to measure the progression of mild cognitive impairment (MCI) and early Alzheimer’s disease (AD). We refer in the following to this first dataset as to the *ADNI* dataset.

A second dataset was generated by Petra Ritter and co-workers at the Charité Hospital in Berlin, with the aim of studying and investigating changes of structural and static and dynamic functional connectivity occurring through healthy aging. This dataset was previously investigated in Zimmermann et al. (2016) and Battaglia et al. (2020) among others. We refer to this second dataset in the following as to the *healthy aging* dataset.

### ADNI dataset

#### Data Sample

Raw neuroimaging data from the Alzheimer’s Disease Neuroimaging Initiative (ADNI) GO/2 studies (Wyman et al., 2013; Beckett et al., 2015) were downloaded for 244 subjects. These included T1w images for all subjects, as well as DWI and rsfMRI images for separate cohorts of subjects. An additional 12 subjects for which both DWI and rsfMRI were acquired in the same session were identified and their data also downloaded.

A volumetric 96-ROI parcellation was defined on the MNI template and consisted of 82 cortical ROIs from the Regional Map parcellation (Kötter & Wanke, 2005) and an additional 14 subcortical ROIs spanning the thalamus and basal ganglia. Details on the construction of the 96-ROI parcellation can be found in Bezgin et al (2017).

Among the 244 subjects we downloaded, 74 were control subjects, while the others were patients at different stages of the pathology progression. In this study, we performed a rough coarse-graining of the original ADNI labels indicating the stage or type of pathology. We thus overall labeled 119 patients as “MCI” (grouping together the labels 4 patients as “MCI”, 64 as “EMCI” and 41 as “LMCI”) and 51 patients as “AD” (overall 170 “Patients” for the simple classification experiments of Figure 6).

Overall, T1 and DTI were jointly available for 88 subjects (allowing to reconstruct structural connectivity (SC) matrix), and T1 and fMRI for 178 (allowing to reconstruct functional connectivity (FC)). However, among the 244 subjects we downloaded, only 12 subjects (referred to as the “SC_emp_+FC_emp_” subset) had a complete set of structural and functional images (T1, DTI, fMRI), hinting at how urgently needed is data completion.

#### Data Preprocessing

Neuroimaging data preprocessing was done using a custom Nipype pipeline implementation (Gorgolewski et al., 2011). First, raw neuroimaging data were reconstructed into NIFTI format using the dcm2nii software package (https://www.nitrc.org/projects/dcm2nii/). Skull stripping was performed using the Brain Extraction Tool (BET) from the FMRIB Software Library package (FSL v5) for all image modalities prior to all other preprocessing steps. Brain extraction of T1w images using BET was generally suboptimal and was supplemented by optiBET (Lutkenhoff et al., 2014), an iterative routine that improved brain extractions substantially by applying transformations and back-projections between the native brain mask and MNI template space. Segmentation of the T1w images was performed using FSL’s FAT tool with bias field correction to obtain into three distinct tissue classes.

To improve the registration of the ROI parcellation to native space, the parcellation was first nonlinearly registered to a publicly-available older adult template (aged 70-74 years, Fillmore et al., 2015) using the Advanced Normalization Tools (ANTS, Avants et al., 2011) software package before subsequent registrations.

Diffusion-weighted images were preprocessed using FSL’s *eddy* and *bedpostx* tools. The ROI parcellation was first nonlinearly registered to each subject’s T1w structural image and then linearly registered to the DWI image using ANTS.

rsfMRI data were preprocessed using FSL’s FEAT toolbox. Preprocessing included motion correction, high-pass filtering, registration, normalization, and spatial smoothing (FWHM: 5 mm). Subjects with excessive motion were excluded from our sample. Global white matter and cerebrospinal fluid signals (but not global mean signal) were linearly regressed from the rsfMRI data.

All images were visually inspected following brain extraction and registrations to ensure correctness.

#### SC Construction

Details of tractography methods for reconstructing each subject’s structural connectome can be found in Shen et al (2019 a, b). Briefly, FSL’s *probtrackx2* was used to perform tractography between all ROIs.

The set of white matter voxels adjacent to a grey matter ROI was defined as the seed mask for that particular ROI. Grey matter voxels adjacent to each seed mask were used to define an exclusion mask. For intra-hemispheric tracking, an additional exclusion mask of the opposite hemisphere was additionally defined. Tractography parameters were set to a curvature threshold of 0.2, 5000 seeds per voxel, a maximum of 2000 steps, and a 0.5 mm step length. The connection weight between each pair of ROIs was computed as the number of streamlines detected between the ROIs, divided by the total number of streamlines sent from the seed mask. This connectivity information was compiled for every subject in a matrix of empirical structural connectivity SC_emp_.

#### rsfMRI Timeseries and FC Construction

Empirical rsfMRI time-series for each ROI were computed using a weighted average approach that favored voxels nearer the center of each ROI (Shen et al., 2012). Each subject’s matrix of empirical functional connectivity FC_emp_ was determined by Pearson correlation of these recorded rsfMRI time-series.

### Healthy aging dataset

#### Data Sample

Forty-nine healthy subjects between the ages of 18 and 80 (mean 42.16 ± 18.37; 19 male/30 female) were recruited as volunteers. Subjects with a self-reported history of neurological, cognitive, or psychiatric conditions were excluded from the experiment. Research was performed in compliance with the Code of Ethics of the World Medical Association (Declaration of Helsinki). Written informed consent was provided by all subjects with an understanding of the study prior to data collection, and was approved by the local ethics committee in accordance with the institutional guidelines at Charité Hospital, Berlin.

#### Acquisition procedures

Acquisition procedures for this data (Magnetic resonance acquisition procedure, dwMRI Data Preprocessing and Tractography, fMRI Data Preprocessing, computation of SC and FC connectome matrices) have been described by Zimmermann et al. (2013), where we redirect the reader interested in full detail.

Briefly, functional and structural image acquisition was performed on a 3T Siemens Tim Trio Scanner MR equipped with a 12-channel Siemens head coil. After anatomical and dwMRI measurements, subjects were removed from the scanner and again put in later for the functional measurements. Data were obtained from subjects at resting state; subjects were asked to close their eyes, relax, and avoid falling asleep.

Anatomical and diffusion images were preprocessed using a fully automated open-source pipeline for extraction of functional and structural connectomes (Schirner et al., 2015). The pipeline performed the following steps. Using the FreeSurfer software toolbox (http://surfer.nmr. mgh.harvard.edu/), anatomical T1-weighted images were motion corrected and intensity normalized, nonbrain tissue was removed, and a brain mask was generated. White matter and subcortical segmentation was performed, and a cortical parcellation based on the probabilistic Desikan– Killiany Freesurfer atlas divided the gray matter into 68 ROIs (regions of interest, 34 per hemisphere) (Desikan et al., 2006). The diffusion data were further corrected (for head movement, eddy current distortions, etc.). Probabilistic fiber tracking was performed using MRTrix streamtrack algorithm.

The fMRI resting-state preprocessing was performed using the FEAT (fMRI Expert Analysis Tool) Version 6.0 first-level analysis software tool from the FMRIB (Functional MRI of the Brain) Software Library (www.fmrib.ox. ac.uk). MCFLIRT motion correction was used to adjust for head movement. Nuisance variables were regressed from the BOLD signal, including the six motion parameters, mean white matter, and CSF signals. Regression of global mean was not performed.

### Two types of computational whole brain models

To bridge between SC and FC via dynamics, we relied on computational modelling of whole-brain intrinsic dynamics. We used two categories of models differing in their complexity, Stochastic Linear Models (SLM) and fully non-linear Mean-Field Models (MFM). SLM procedures are used for linear SC-to-FC and FC-to-SC completions, while MFM procedures are used for analogous but nonlinear completions.

### SLM models

The SLM model used in this study is a linear stochastic system of coupled Ornstein-Uhlenbeck processes which is deeply investigated in (Saggio et al., 2016). For each brain region, neural activity *x*_*i*_(*t*) is modeled as a linear stochastic model, coupled to the fluctuations of other regions:

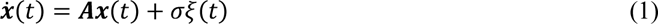

where ***A*** is the coupling matrix, *ξ* is a normal Gaussian white noise, and *σ* the standard deviation of the local drive noise. The coupling matrix ***A*** can be written as:

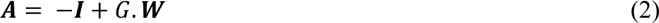

where ***I*** is the identity matrix, *G* is the global coupling parameter and ***W*** is a weight matrix set to match SC_emp_. The negative identity matrix guarantees that the nodes have a stable equilibrium point. If all the eigenvalues of ***A*** are negative, which happens for all positive values of *G* < *G*_*critic*_ = 1/*max*(*λ*) (where *λ*_*i*_ are the eigenvalues of ***W***), the system will be in an equilibrium state. After some mathematical steps (Saggio et al., 2016), the covariance matrix between regional fluctuations can be analytically expressed at this critical point *G_critic_* as:

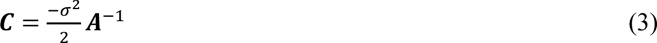

whose normalized entries provide the strength of functional connectivity between different regions. The noise strength can be arbitrarily set at the critical point since it provides only a scaling constant to be reabsorbed into the Pearson correlation normalization. However, the only parameter that needs to be explored is *G*, whose range goes from *G_min_* = 0, i.e. uncoupled nodes, to slightly before *G_critic_* = 1/*max*(*λ*_*i*_), or *G_max_* = *G_critic_* – *ε*. In Extended Data Figure 3-1A, running explicit simulations of SLM models for different values of coupling *G* and evaluating on the “FC_emp_ + SC_emp_” subset of ADNI subjects the match between the simulated and empirical activity correlation matrices, we confirm (cf. e.g. Hansen et al., 2015) that the best match (max of Pearson correlation between the upper-triangular parts of the empirical and virtual FCs) is obtained at a slightly subcritical point for *G\** = *G_critic_* – *ε*.

### Linear SC-to-FC and FC-to-SC completion

To infer FC_SLM_ from SC_emp_, we chose to always use a common value *G*_ref_* = 0.83, which is the median of *G\** for all 12 “FC_emp_ + SC_emp_” subjects in the ADNI and Healthy Ageing dataset (the error made in doing this approximation is estimated to be less than 1% in Extended Data Fig. 3-1 C). When the connectome FC_emp_ is not known, equations (2) and (3) can directly be used to evaluate the covariance matrix ***C*** (setting σ = 1 and *G = G*_ref_*). We then estimate the regional fluctuation covariance from these inferences and normalize it into a Pearson correlation matrix to infer FC_SLM_ (*See pseudo-code in Table 1-1*). Linear FC_SLM_ completions for our ADNI dataset and for the Healthy Aging dataset can be downloaded as MATLAB® workspace within Extended Data FC_SLM.mat (available at the address https://github.com/FunDyn/VirtualCohorts).

To infer SC_SLM_ from FC_emp_, we invert the analytical expressions of eqs. (2) and (3) and always set *σ* = 1 and

*G = G*_ref_* leading to:

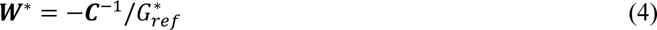

where ***C*** is the covariance matrix estimated from empirical BOLD time-series. The linearly completed SC_SLM_ is then set to be identical to ***W****\** setting its diagonal to zero to avoid offsets, which would be meaningless given the conventional choice of noise σ which we have made (*see Table 2-1*). Note that all the free parameters of the SLM model appear uniquely as scaling factors and do not affect the (normalized) correlation of the inferred SC_SLM_ with the SC_emp_. However, the absolute strengths of inferred structural connections remain arbitrary, with only the relative strengths between different connections being reliable (since unaffected by arbitrary choices of scaling parameters; *see pseudo-code in Table 2-1*). Linear SC_SLM_ completions for the ADNI dataset and for the Healthy Aging dataset can be downloaded as MATLAB® workspace within Extended Data SC_SLM.mat (available at the address https://github.com/FunDyn/VirtualCohorts).

### MFM models

For non-linear completion algorithms, we performed simulations of whole-brain mean-field models analogous to Deco et al. (2013) or Hansen et al. (2015). We used a modified version of the mean-field model designed by Wong and Wang (2006), to describe the mean neural activity for each brain region, following the reduction performed in (Deco et al., 2013). The resulting neural mass equations are given by:

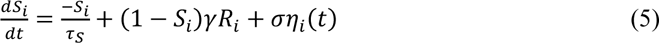

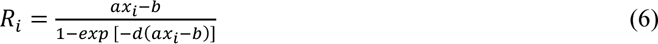

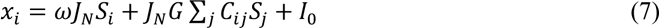

where *τ*_*i*_ represents NMDA synaptic input currents and *τ*_*s*_ the NMDA decay time constant; *R*_*i*_ is collective firing rates; γ = 0.641 is a kinetic parameter; *a* = 270(*V*. *nC*)^-1^, *b* = 108*Hz*, *d* = 0.154*s* are parameters values for the input-output function; *x*_*i*_are the total synaptic inputs to a regions; *J*_*N*_ = 0.2609*nA* is an intensity scale for synaptic currents; *ω* is the relative strength of recurrent connections within the region; *C*_*ij*_ are the entries of the SC_emp_ matrix reweighted by global scale of long-range connectivity strength *G* as a control parameter; *σ* is the noise amplitude, and *η*_*i*_ is a stochastic Gaussian variable with a zero mean and unit variance. Finally, *I*_0_ represents the external input and sets the level of regional excitability. Different sets of parameters yield different neural network dynamics and, therefore, patterns of FC_MFM_ non-stationarity.

To emulate BOLD fMRI signals, we then transformed the raw model output activity *x*_*i*_ through a standard Balloon-Windkessel hemodynamic model. All details of the hemodynamic model are set according to Friston et al. (2003).

### Non-linear SC-to-FC completion

In general, our simple MFM model has three free parameters at the level of the local neural mass dynamics (*τ, ω*, and *I_0_*) and one free global parameter *G.* Since changing the values of *ω* and *I_0_* had lesser effects on the collective dynamics of the system (see Extended Data Figure 3-2), we set their values to *ω* = 0.9 and *I_0_ =* 0.32 respectively and remain then just two free parameters which we allow to vary in the ranges *G* ∈ [1 3] and *τ* ∈ [1 100] *ms* when seeking for an optimal working point of the model. As revealed by the analyses of Figure 3, the zone in this restricted parameter space associated with the best FC-rendering performance can be identified through the joint inspection of three scores, varying as a function of both *G* and *τ*. The first criterion is the spatial heterogeneity of activation (*see Table 1, line 2.5*) computed by taking the coefficient of variation of BOLD_MFM_ time-series.

**Figure 1.**
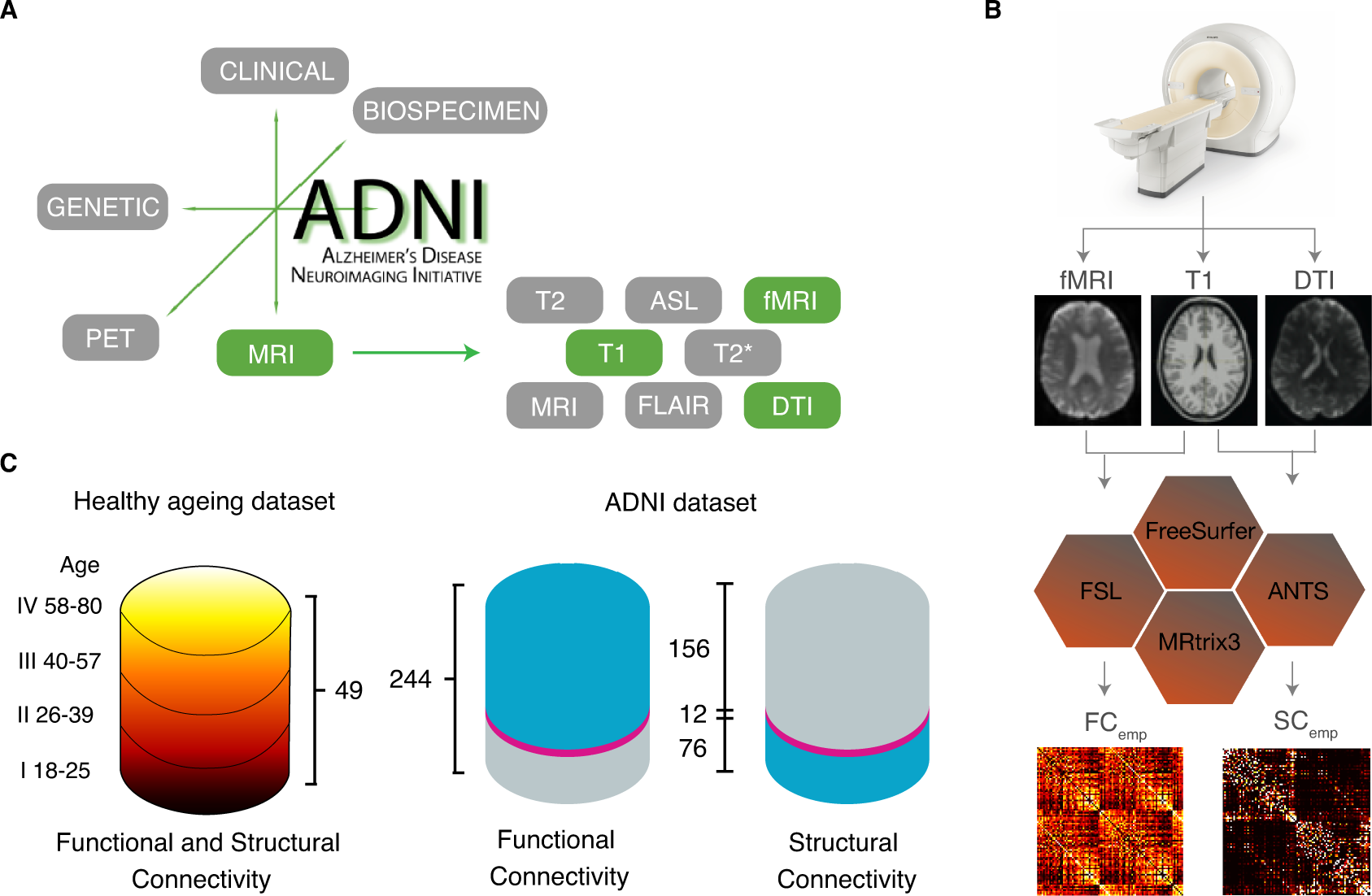
Connectomic information extracted from the ADNI dataset has gaps. A) The different dataset releases by the ADNI consortium include a variety of information relative to different biomarkers and imaging modalities. Here, we focus on structural and functional MRI features and, chiefly: T1, DTI (allowing to extract empirical structural connectomes); and resting-state fMRI BOLD time-series (allowing to extract empirical functional connectomes). B) Matrices SC_emp_ and FC_emp_ summarizing connectomic information about, respectively structural connectivity (SC) and functional connectivity (FC) are obtained via elaborated multi-step processing pipelines, using various software including FreeSurfer, FSL, ANTS, and MRtrix3. C) The total number of subjects in Healthy ageing dataset is 49 between the ages of 18 and 80 (mean = 42.16 ± 18.37; 19 male/30 female) in which with approximately equal number of subjects they were divided into 4 categories (I:IV). The total number of ADNI-derived subjects investigated in this study is 244, in which 74 subjects were control, while 119 subjects labeled as MCI, and 51 subjects as AD. Out of these 244, FC_emp_ could be extracted for 168 subjects, and SC_emp_ for 88. However, SC_emp_ and FC_emp_ were both simultaneously available for just a minority of 12 subjects (referred to as the “SC_emp_+FC_emp_ subset”). The available data is shown in blue and the missing data in grey, the SC_emp_+FC_emp_ subset is shown in pink.

**Figure 2.**
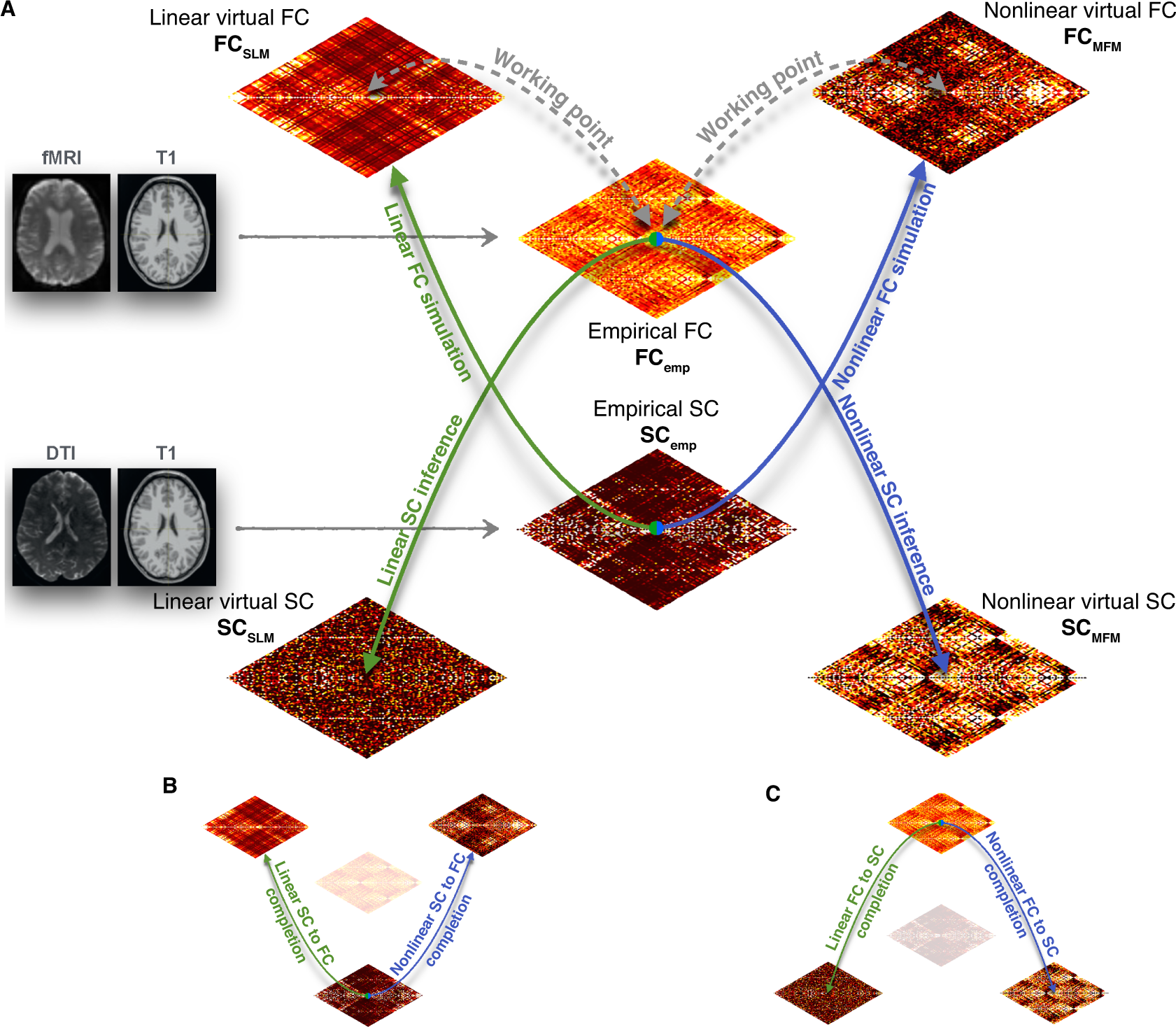
From mean-field modeling to connectomic data completion. A) We present here a graphical summary of the various computational simulation and inference strategies used in this study to bridge between different types of connectivity matrices. Mean-field simulation and the associated analytic theory can be used to generate virtual FC, through simulations of resting-state whole-brain models embedding a given input SC connectome (ascending arrows). Algorithmic procedures, that may still include computational simulation steps, can be used to perform the inverse inference of a virtual SC that is compatible with a given input FC (descending arrows). Both simulation and inference can be performed using simpler linear (green arrows) or non-linear (blue arrows) approaches. When the input SC (or FC) connectomes used as input for FC simulation (or SC inverse inference) correspond to empirical connectomes SC_emp_ (or FC_emp_), derived from T1 and DTI (fMRI) images, then model simulation (inversion) can be used to complete gaps in the dataset, whenever FC_emp_ (or SC_emp_) is missing. We refer then to these operations as: B) SC-to-FC completion; and, C) FC-to-SC completion. Both exist in linear and non-linear versions.

**Figure 3.**
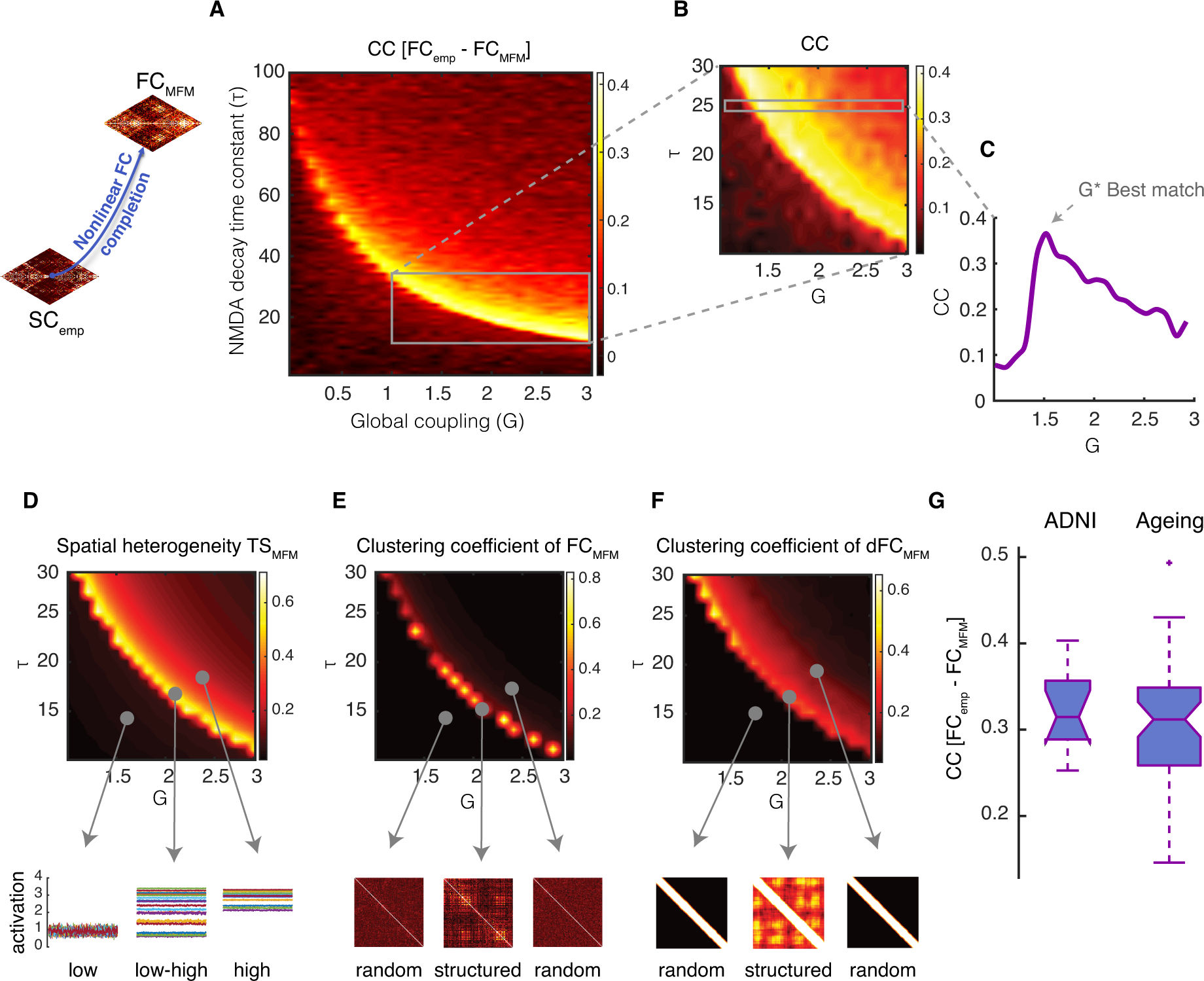
Non-linear SC-to-FC data completion. Simulations of a non-linear model embedding a given input SC_emp_ matrix can be used to generate surrogate FC_MFM_ matrices. A) Systematic exploration (here shown for a representative subject) of the dependency of the correlation between FC_emp_ and FC_MFM_ on the MFM parameters *G* (inter-regional coupling strength) and *τ* (synaptic time-constant of within-region excitation) indicates that the best fitting performances are obtained when parameters are concentrated in a narrow concave stripe across the *G/τ* plane. B) Enlarged zoom of panel A over the range *G* ∈ [1 3] and *τ* ∈ [10 30]. C) For a value of *τ* = 25, representatively chosen here for illustration, we identify a value *G\** for which the Pearson correlation between FC_emp_ and FC_MFM_ reaches a clear local maximum. Panels A-C thus indicate that it makes sense speaking of a best-fit zone and that reliable nonlinear SC-to-FC completion should be performed using MFM parameters within this zone. Three criteria help us identifying parameter combinations in this best fitting zone when the actual FC_emp_ is unknown. D) First criterion: we define the spatial coefficient of variation of the time-series of simulated BOLD activity TS_MFM_ as the ratio between the variance and the mean across regions of the time-averaged activation of different regions. The best fit zone is associated with a peaking of this spatial coefficient of variation, associated with a maximally heterogeneous mix or low and high activation levels for different regions (see time-series in lower cartoons). E) Second criterion: in the best fitting zone, the resulting FC_MFM_ is neither randomly organized nor excessively regular (synchronized) but presents a complex clustering structure (see lower cartoons), which can be tracked by a peak in the clustering coefficient of the FC_MFM_, seen as weighted adjacency matrix. F) Third criterion: in the best fitting zone, resting-state FC_MFM_ display a relatively richer dynamics than in other sectors of the parameter space. This gives rise to an “dFC matrix” (correlation between time-resolved FC observed at different times) which is neither random nor too regular but displays a certain degree of clustering (see lower cartoons). The emergence of complex dynamics of FC can be tracked by an increase in the clustering coefficient of the dFC matrix extracted from simulated resting-state dynamics. G) The boxplot shows the distribution of correlations between the actual FC_emp_ and FC_MFM_ estimated within the best fitting zone for all subjects from the “SC_emp_ + FC_emp_” ADNI subset and the ageing dataset. See Extended Data Figure 3-1 for linear SC-to-FC completion and Extended Figure 3-2 for dependency of MFM best fit zone on additional parameters.

**Figure 4.**
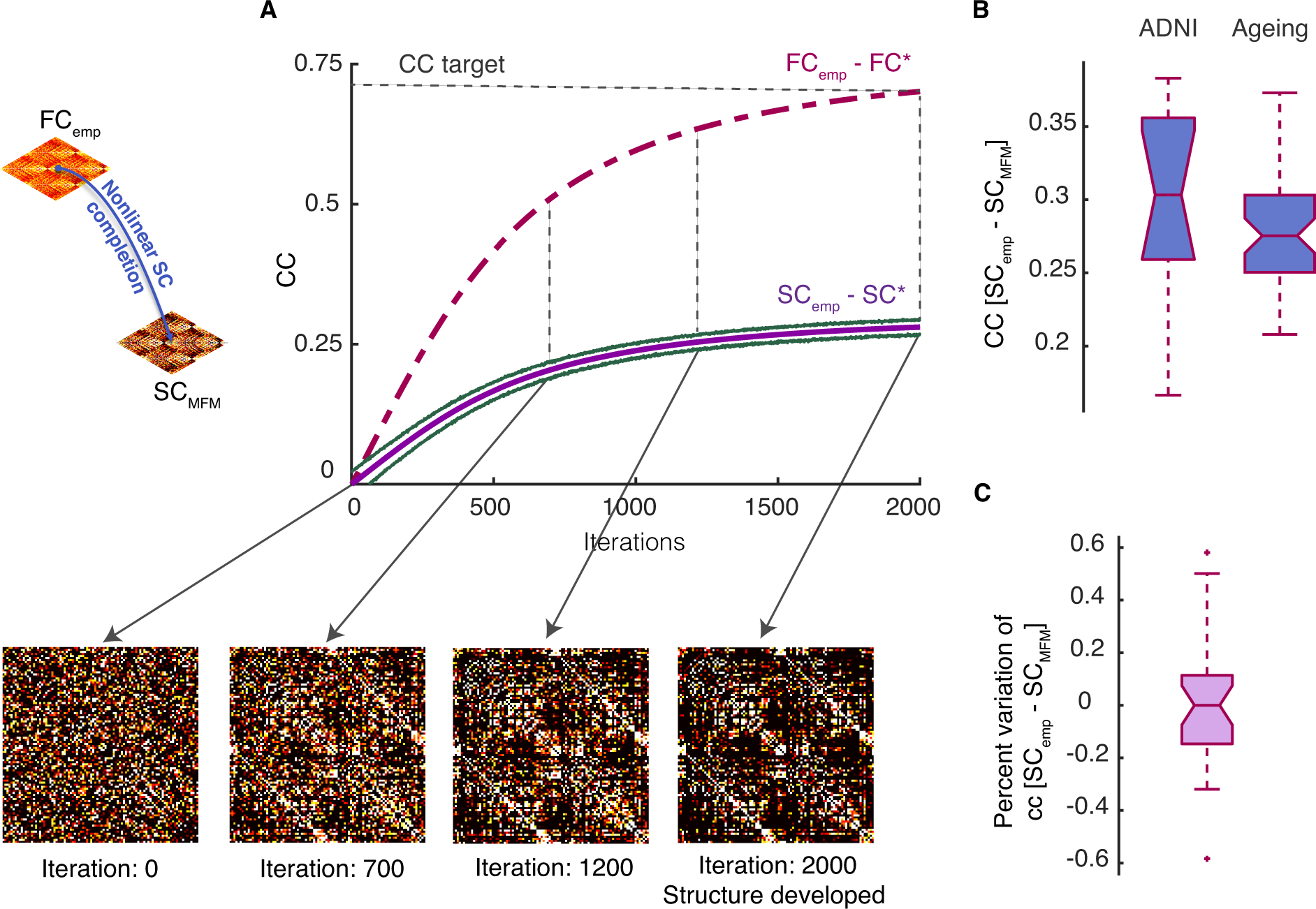
Non-linear FC-to-SC data completion. An iterative procedure can be used to perform resting-state simulations of an MFM model starting from a randomly guessed structural connectome SC* and progressively modify this SC* to make it compatible with a known target FC_emp_. A) Starting from an initial random SC*_(0)_ matrix, there is no correlation between the target FC_emp_ and the generated FC*_(0)_ matrix. However, by adjusting the weights of the used SC* through the algorithm of Table 2, SC* gradually develops a richer organization, leading to an increase of the correlation between FC* and FC_emp_ (violet dashed line) and in parallel, of the correlation between SC* and SC_emp_ (violet solid line), as shown here for a representative subject within the “SC_emp_+FC_emp_” subset. The algorithm stops when the correlation between FC* and the input target FC_emp_ reaches a desired quality threshold (here 0.7 after 2000 iterations) and the SC* at the last iteration is used as virtual surrogate SC_MFM_. B) The boxplot shows the distribution of correlation between SC_emp_ and SC_MFM_ for all subjects in the “SC_emp_ + FC_emp_” ADNI subset and the Healthy Ageing dataset. C) The correlation between SC_emp_ and SC_MFM_ can vary using different random initial connectomes SC*_(0)_. Here we show a boxplot of the percent dispersions of the correlation values obtained for different initial conditions around the median correlation value. The fact that these dispersions lie within a narrow interval of ±2.5% indicates that the expected performance is robust against changes of the initial conditions. See Extended Data Figure 4-1 for linear FC-to-SC completion.

**Table 1.**
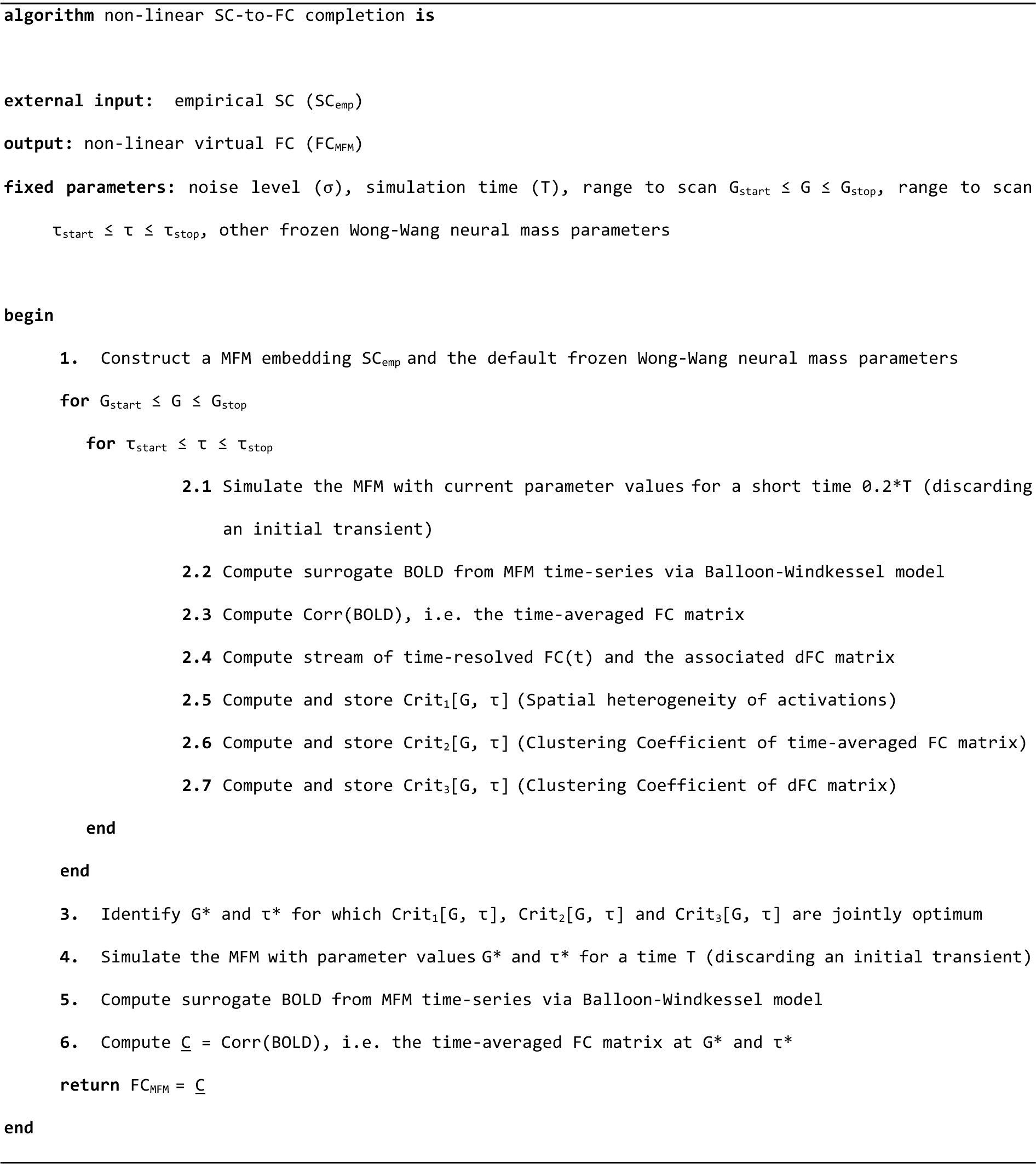
Pseudo-code for non-linear SC-to-FC completion (FC virtual duals to SC)

By computing the Pearson correlation coefficient of upper-triangular between FC_MFM_ and FC_emp_ for every subject from “SC_emp_ + FC_emp_” subset in the ADNI dataset (*see Table 1, line 2.3*), we obtained a best-fitting zone in a narrow concave stripe (see Figure 3A for one subject); (*G*^*^, *τ*^*^) parameter set, bring the system to this best-fitting zone and values lower than this is (*G*^”^, *τ*^”^) set and higher values are (*G*^+^, *τ*^+^). Qualitatively analogous results are found for the healthy aging dataset. This non-monotonic behavior of yellow zone in *G/τ* plane occurs where three criteria are jointly met; the second criterion is the clustering coefficient of time-average FC_MFM_ matrices (*see Table 1, line 2.6*) and finally, the third criterion is the clustering coefficient of dFC_MFM_ matrices (*see Table 1, line 2.6*), where the dFC matrices were computed for an arbitrary window using the dFCwalk toolbox (Arbabyazd et al., 2020; https://github.com/FunDyn/dFCwalk.git). By knowing the optimal working point of the system where all three criteria are jointly optimum (*see Table 1, line 2*), we freeze the algorithm and finally run a last simulation with the chosen parameters to perform non-linear SC-to-FC data completion (*see Table 1, lines 3 to 5*). Non-linear FC_MFM_ completions for our ADNI dataset and for the Healthy Aging dataset can be downloaded as a MATLAB® workspace within Extended Data FC_MFM.mat (available at the address https://github.com/FunDyn/VirtualCohorts).

### Non-linear FC-to-SC completion

We implemented a heuristic approach to infer the most likely connectivity matrix (i.e. Effective Connectivity) that maximizes the similarity between empirical and simulated functional connectivity. As an initial point, we considered a random symmetric matrix and removed diagonal as SC*_(0)_ (*see Table 2, line 1*) and run the algorithm in Table 1 in order to simulate the FC*_(0)_. Then iteratively we adjusted the SC as a function of the difference between the current FC and empirical FC (*see Table 2, line 2*), in other words SC*_(1)_ = SC*_(0)_ + λΔFC_(0)_ where ΔFC_(0)_ = FC_emp_ – FC*_(0)_ and λ is the learning rate (*see Table 2, line 3*). The iteration will stop when the correlation between FC_emp_ and FC*_(k)_ reaches to the threshold CC_target_ = 0.7 and giving the SC*_(k)_ as SC_MFM_. All the parameter used in this section is identical to the non-linear SC-to-FC completion procedure. Nonlinear SC_MFM_ completions for our ADNI and healthy aging datasets can be downloaded as a MATLAB® workspace within Extended Data SC_MFM.mat (available at the address https://github.com/FunDyn/VirtualCohorts).

**Table 2.**
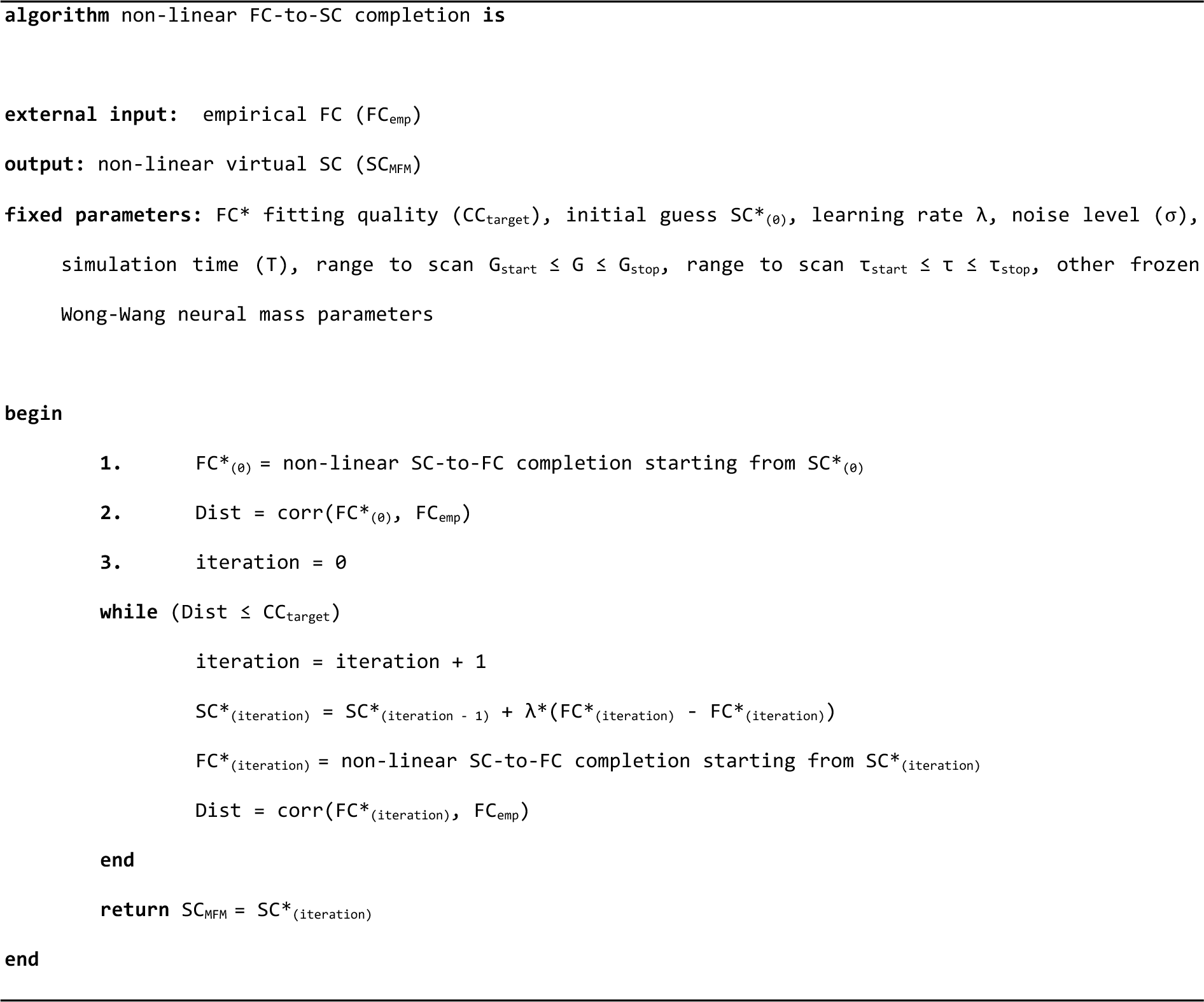
Pseudo-code for non-linear FC-to-SC completion (SC virtual duals to FC)

### Trivial completion using the “other connectome”

In the case in which one of the two connectomes is missing (e.g. just SC available but not FC) one may think to use the available connectome (in this example, SC) as a “good guess” for the missing one (in this example, FC). We refer to this trivial procedure as a completion using the other connectome. If the match quality between surrogate connectomes obtained via more complex procedures and the target empirical connectome to reconstruct happened to be comparable with the one that one can get via the trivial completion, then it would not be worth using more sophisticated methods. We assessed then, for comparison with other strategies, the performance of such trivial completion approach on the “SC_emp_ + FC_emp_” subset of the ADNI dataset and on the whole Healthy Aging dataset. In order for a completion approach to be considered viable, it is necessary that it outperforms significantly this trivial completion via the “other type” connectome, which can be quantified by a relative improvement coefficient:

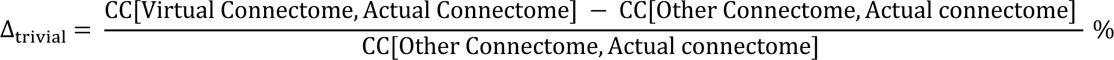

### Bi-virtual data completion

The pipelines for data completion described above can be concatenated, by performing e.g. FC-to-SC completion on a virtually FC or SC-to-FC completion on a virtual SC (rather than actual FC_emp_ or SC_emp_, respectively). In this way, one can create bi-virtual dual counterparts SC_bi-MFM_ (FC_bi-MFM_) or SC_bi-SLM_ (FC _bi-SLM_) for any of the available empirical SC_emp_ (FC_emp_) by applying in sequence non-linear MFM-based or linear SLM-based procedures for SC-to-FC and then FC-to-SC completion (or, conversely, FC-to SC followed by SC-to-FC completions). Linear and nonlinear bi-virtual completions for our ADNI and Healthy Aging datasets can be downloaded as MATLAB® workspaces within Extended Data SC_bivirt.mat and FC_bivirt.mat (available at the address https://github.com/FunDyn/VirtualCohorts).

For every pair of subjects, we computed the correlation distance between the respective empirical connectomes (pairs of FC_emp_ or SC_emp_) and the corresponding bivirtual duals (pairs of FC_bi-MFM_ or SC_bi-MFM_) and plotted the empirical-empirical distances vs the corresponding bivirtual-bivirtual distances (cf. Figure 6) to reveal the large degree of metric correspondence between real and bivirtual dual spaces. This correspondence was also quantified computing Pearson Correlation between empirical and bivirtual pairwise distances. These correlations (computed as well for virtual connectomes, beyond the bivirtual duals) are tabulated in Table 4.

**Figure 5.**
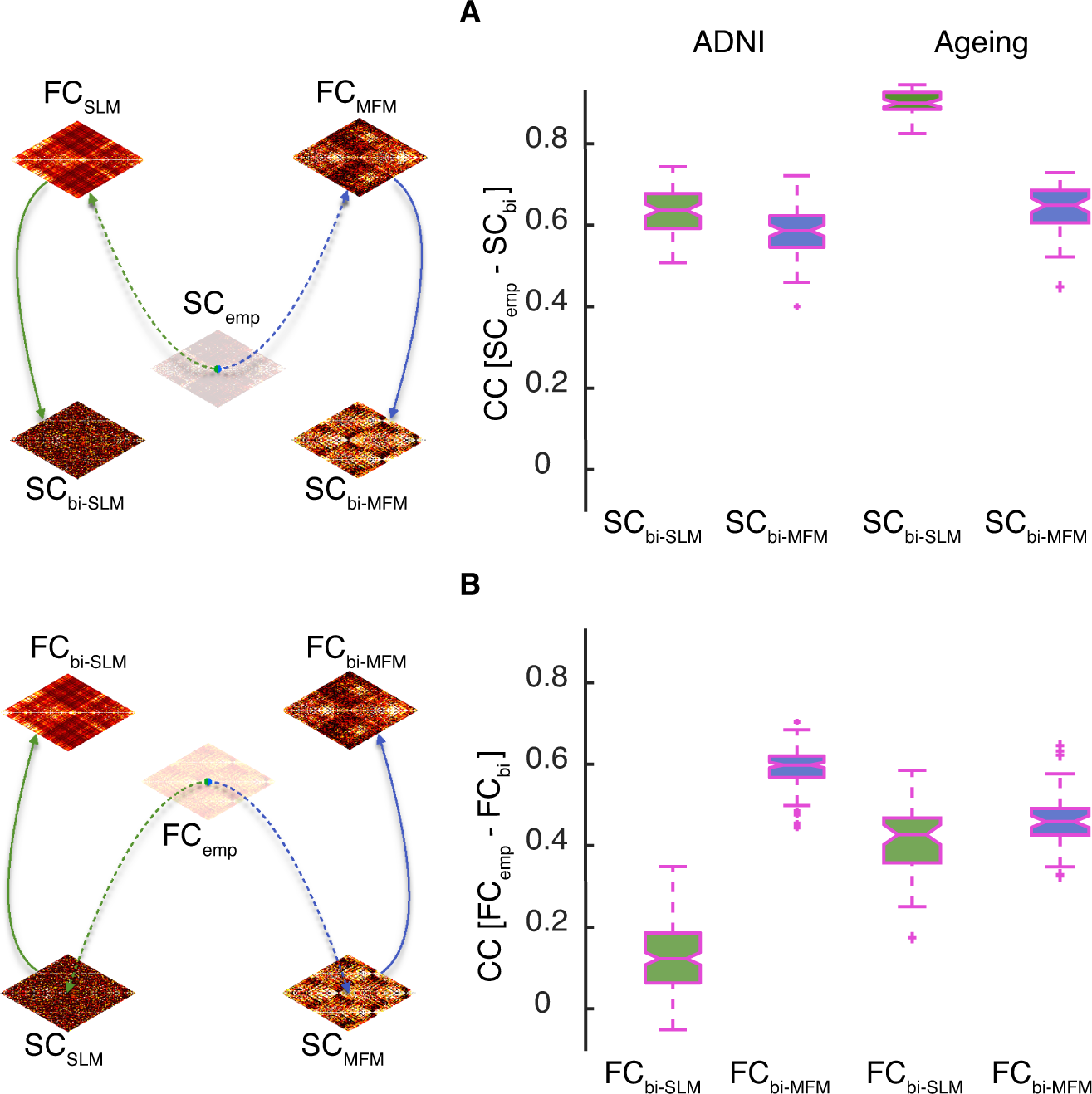
Bi-virtual connectomes. This figure shows the correspondence between empirical and bi-virtual SC and FC pairs, both when using chained linear (SLM-based) and nonlinear (MFM-based) completion procedures. A) For 88 subjects from the ADNI-subset with only SC_emp_ available, considering the linear bi-virtual completion chain SC_emp_ to FC_SLM_ to SC_bi-SLM_, we obtained a median correlation between SC_emp_ and SC_bi-SLM_ equal to 0.63 and 0.92 for 49 subjects from the Healthy Ageing dataset (green boxplot); simultaneously, considering the non-linear bi-virtual completion chain SC_emp_ to FC_MFM_ to SC_bi-MFM_, we obtained a median correlation between SC_emp_ and SC_bi-MFM_ equal to 0.58 for the ADNI datast and 0.64 for the Healthy Ageing dataset (blue boxplot). B) For 168 subjects from the ADNI-subset with only FC_emp_ available, considering the linear bi-virtual completion chain FC_emp_ to SC_SLM_ to FC_bi-SLM_, we obtained a median correlation between FC_emp_ and FC_bi-SLM_ equal to 0.12 and 0.42 for 49 subjects from Healthy Ageing dataset (green boxplot); simultaneously, considering the non-linear bi-virtual completion chain FC_emp_ to SC_MFM_ to FC_bi-MFM_, we obtained a median correlation between FC_emp_ and FC_bi-MFM_ equal to 0.59 for the ADNI dataset and 0.45 for the Healthy Ageing dataset (blue boxplot).

**Figure 6.**
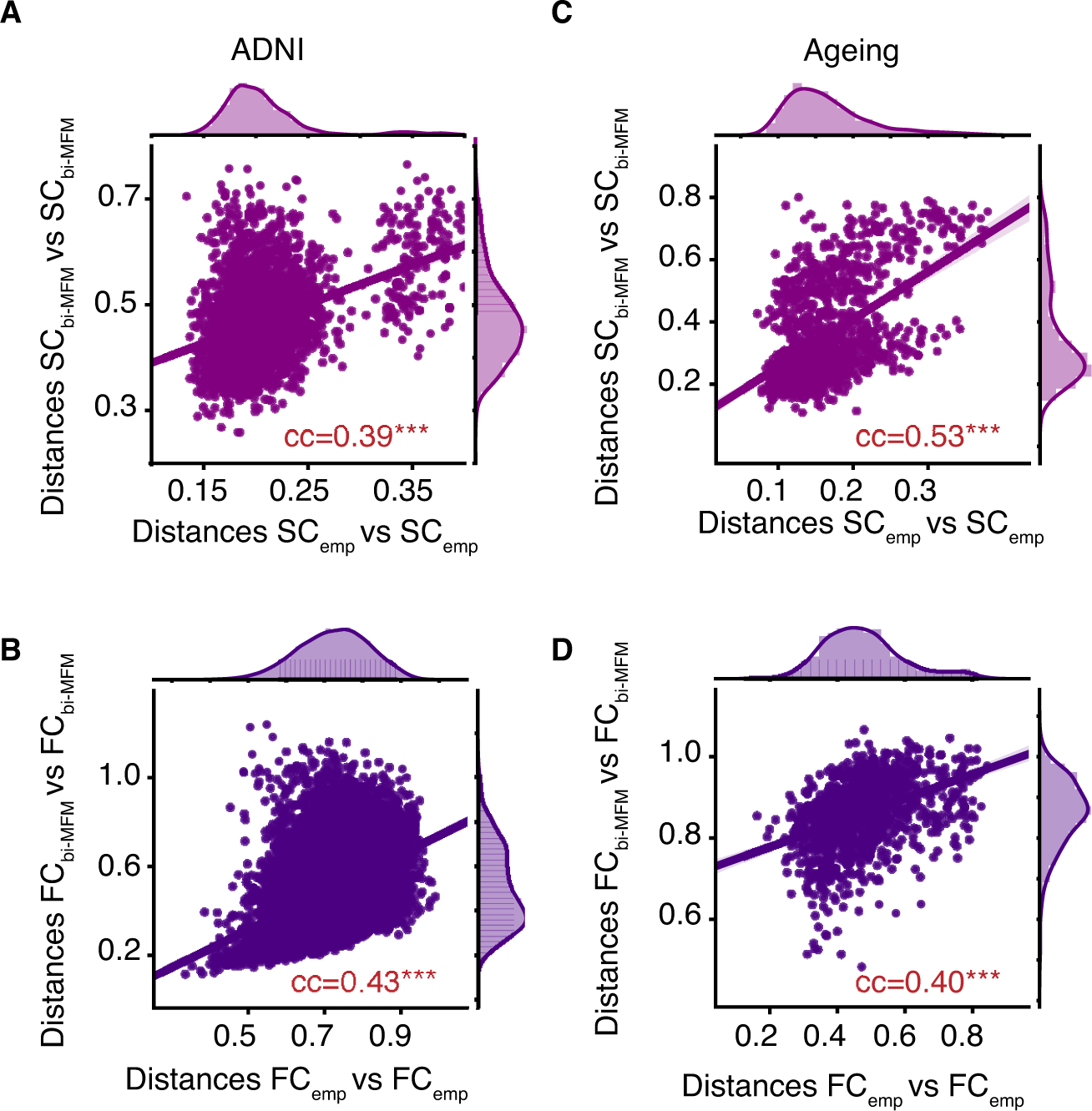
Inter-subject distances for empirical – bivirtual pairs. We show here the distances between the empirical SC_emp_ (or FC_emp_) of different subjects and the inter-subject distances for their corresponding pairs of subjects from bivirtual SC_bi-MFM_ (or FC_bi-MFM_). A-B) For the ADNI dataset the correlation between the inter-subject distances in real and dual spaces for SC (between SC_emp_ and SC_bi-MFM_) were significant and equal to 0.39, and for FC pairs (between FC_emp_ and FC_bi-MFM_) equal to 0.43. C-D) The same inter-subject distances for the healthy ageing dataset were measured, with correlation values equal to 0.53 and 0.40 for SC and FC empirical-bivirtual pairs, respectively.

### Improvement by personalization

Completion procedures map a connectome for a given subject to subject-specific virtual and bivirtual dual connectomes. The question is whether the similarity between empirical and completed connectomes is better when considering connectome pairs formed by an empirical and its subject-specific dual connectomes, or pairs made by an empirical and a generic virtual or bivirtual connectome, not specific to the considered subject. We expect that empirical-to-virtual match is improved by personalization. To quantify it, we introduce an Improvement by Personalization coefficient Δ_Pers_, evaluating it for all the types of completion.

For simulated data one can define CC_personalized_ = CC[Connectome_virt_(a subject), Connectome_emp_(same subject)], where “Connectome” refers to the considered connectome matrix (of either the SC or the FC type) and the ondex “virt” to any type of completion (SLM- or MFM-based, virtual or bivirtual). Analogously, we define CC_generic_ = Group average of CC[Connectome_virt_(same subject), Connectome_emp_(a different subject)]. The Improvement by Personalization coefficient is then defined as Δ_Pers_ = (CC_personalized_ - CC_generic_) / CC_generic_. This coefficient significantly larger than zero denotes that completion pipelines get to improved results when completion is personalized.

At least for Functional Connectivity, we can estimate from empirical data how much the improvement by personalization could be expected to be in the case in which a first FC extraction for a given subject had to be replaced by a second one coming from a second scan from the same subject vs a scan for another generic subject. To obtain such an estimate, we focus on a dataset mediated from the Human Connectome Project and conceived to probe test/retest variability (Termenon et al., 2016). In this dataset, 100 subjects underwent two resting state scans, so that two FC_emp_ can be extracted for each of them. If we redefine CC_personalized_ = CC[FC_emp_(same subject first scan), FC_emp_(same subject second scan)] and CC_generic_ = Group average of CC[FC_emp_(same subject, first scan), FC_emp_(a different subject, first scan)], then we can evaluate an empirical Δ_Pers_ = (CC_personalized_ - CC_generic_) / CC_generic_. For empirical FCs from the Termenon et al. (2016) dataset we obtain an improvement by personalization of ∼+22%, to be used as a comparison level when looking at improvements by personalization in virtual and bivirtual connectomes.

### Network topology features and their personalized preservation through data completion

To evaluate the correspondence between empirical and bivirtual connectomes we evaluated a variety of graph-theoretical descriptors of the connectomes and compared them within pairs of empirical and bivirtual dual adjacency matrices. Every connectome, functional or structural, was described by a weighted undirected matrix *C_ij_*, where *i* and *j* are two brain regions, and the matrix entries denote the strength of coupling –anatomical or at the level of activity correlations– between them. For each brain region *i*, we then computed: its *strength S_i_ =* Σ*_j_ C_ij_*, indicating how strongly a given region is connected to its local neighborhood; its *clustering coefficient* Clu*_i_ =* |triangles involving *i*| / |pairs of neighbors of *i|* (with |⋅| denoting the count of a type of object), determining how densely connected are between them the neighbors of the considered region; and its *centrality coefficient*, quantifying the tendency for paths interconnecting any two nodes in the networks to pass through the considered node. In particular, we computed here centrality using a version of the *PageRank* algorithm (Brin and Page, 1998) for weighted undirected networks in an implementation from the Brain Connectivity Toolbox (Bullmore & Sporns, 2009), with a typical damping parameter of 0.9. Without entering in the details of the algorithm (see Brin and Page, 1998 for details), a node is deemed important according to PageRank centrality if it receives strong links from other important nodes sending selective and parsimonious in their connections, i.e. sending only a few strong links. Strengths, clustering, and centrality measures provide together a rich and detailed portrait of complementary aspects of network topology and on how it varies across brain regions. We computed then the correlations between the above graph theoretical features for matching regions in empirical connectomes and their bivirtual counterparts. Note that the number of network nodes were different for connectomes in the ADNI and in the healthy aging datasets, since the used reference parcellations included a different number of regions in the two cases. However, graph theoretical metrics can be computed in precisely the same way and we perform in this study uniquely within-dataset analyses. In Figure 8 we show point clouds for all subjects of the ADNI dataset pooled together. Analogous plots for the healthy aging dataset are shown in Figure 8-1.

We then computed correlations between vectors of graph-theoretical features over the different brain regions *within* specific subjects. This analysis is an important probe of the personalization quality in data completion, since every subject may have a different spectrum of graph-theoretical properties across the different regions and that it is important that information about these topological specificities is maintained by completion. These within-subject correlations –often higher than global population correlations, since not disturbed by variations of mean feature values across subjects– are summarized in Table 3 for the ADNI dataset and in Table 3-1 for the healthy aging dataset. In these tables, we provide both absolute correlation values and the indication of how each correlation is improved by computing it within subjects rather than across the whole sample. Correlations were evaluated over data points belonging to the interquartile range of empirical data and then extrapolated to the whole range to avoid estimation to be fully dominated by cloud tails of extreme outliers.

**Table 3.** Single-subject correlations between network features in real and bivirtual dual connectomes for the ADNI dataset

|  | SC |  | FC |  |
| --- | --- | --- | --- | --- |
| | Median and range<br>Within subject<br>*cross-subject* | $\Delta\%$ | Median and range<br>Within subject<br>*cross-subject* | $\Delta\%$ |
| **Strength** | $0.16 \pm 0.20$<br>$0.13 \pm 0.17$ | **$25 \pm 18$** | $0.77 \pm 0.18$<br>$0.17 \pm 0.20$ | **$342 \pm 8$** |
| **Clustering** | $-0.05 \pm 0.12$<br>$-0.06 \pm 0.11$ | $-17 \pm 24$ | $0.65 \pm 0.24$<br>$0.14 \pm 0.21$ | **$359 \pm 13$** |
| **Centrality** | $0.21 \pm 0.18$<br>$0.16 \pm 0.15$ | **$24 \pm 12$** | $0.66 \pm 0.20$<br>$0.16 \pm 0.18$ | **$312 \pm 10$** |
| **Communities** | $59\% \pm 10\%$<br>$47\% \pm 8\%$ | **$23 \pm 2$** | $45\% \pm 10\%$<br>$12\% \pm 6\%$ | **$260 \pm 6$** |

*Indicated values for real/bivirtual dual correlations (for strength, clustering and centrality coefficients) or relative mutual information (for communities) are mean ± standard deviation of the mean over subjects.*

**Table 4.** Single-subject correlations between network features in real and bivirtual dual connectomes for the healthy ageing dataset

|  | SC |  | FC |  |
| --- | --- | --- | --- | --- |
| | Median and range<br>Within subject<br>*cross-subject* | $\Delta\%$ | Median and range<br>Within subject<br>*cross-subject* | $\Delta\%$ |
| **Strength** | $0.80 \pm 0.04$<br>$0.76 \pm 0.07$ | $5 \pm 1$ | $0.65 \pm 0.18$<br>$0.37 \pm 0.16$ | $75 \pm 7$ |
| **Clustering** | $0.83 \pm 0.06$<br>$0.79 \pm 0.08$ | $6 \pm 1$ | $0.64 \pm 0.22$<br>$0.38 \pm 0.19$ | $70 \pm 8$ |
| **Centrality** | $0.80 \pm 0.05$<br>$0.76 \pm 0.06$ | $4 \pm 1$ | $0.63 \pm 0.18$<br>$0.38 \pm 0.16$ | $65 \pm 7$ |
| **Communities** | $44\% \pm 8\%$<br>$38\% \pm 8\%$ | $16 \pm 3$ | $53\% \pm 10\%$<br>$48\% \pm 12\%$ | $10 \pm 3$ |

*Indicated values for real/bivirtual dual correlations (for strength, clustering and centrality coefficients) or relative mutual information (for communities) are mean ± standard deviation of the mean over subjects..*

We extracted then the community structure of empirical and bivirtual dual connectomes using the Louvain algorithm (Blondel et al., 2008), with default parameter Γ = 1 and “negative symmetric” treatment of negative matrix entries (once again, in the implementation of the Brain Connectivity Toolbox). To compare the resulting community assignments to different regions across pairs of dual empirical and bivirtual connectomes we computed the Mutual Information between the respective labelings and normalized it in the unit range by dividing it by the largest among the entropies of the community labelings of each connectome. Such normalised mutual information measure is not sensitive to changes in names of the labels and can be applied independently on the number of retrieved communities. Chance levels for relative mutual information can be estimated by permuting randomly the labels and finding the 99_th_ percentile of values for shuffled labels. Average Mutual Information between community labels are tabulated as well in Table 3 for the ADNI dataset and in Table 3-1 for the healthy aging dataset, once again giving absolute values and relative improvements of personalized with respect to generic correspondence.

### Supervised subject classification

To show the possibility to extract personalized information relevant for subject characterization, we performed different machine-learning supervised classification tasks using as input features derived from empirical and (bi)virtual connectomes. The input and target features to predict were different for the ADNI and the healthy aging datasets.

Concerning the ADNI dataset, we separated subjects in two subgroups: “controls” and “patients” (“MCI” or “AD”). Subjects (the actual ones or their associated virtual counterparts) are thus labeled as “positive” when belonging to the patient subgroup or “negative” otherwise. Note that our classifiers were not sufficiently powerful to reliably discriminate subjects in three classes (“control”, “MCI” and “AD”) on this dataset, at least under the simple classification strategies we used. For illustration, we constructed classifiers predicting subject category from input vectors compiling the total connectivity strengths (in either SC or FC connectomes, real, virtual, or bivirtual) of different brain regions. The dimension of the input space was thus limited to the number of regions in the used 96-ROIs parcellation, which is of the same order of the number of available subjects in the overall dataset.

Concerning the healthy aging dataset, we separated subjects in four age classes with 13 subjects in class I (age = 18-25), and 12 subjects in classes II (age = 26-39), III (age = 40-57), and IV (age = 58-80) and used as target labels for classification the ordinal of the specific age class of each subject. As input vectors we used in this case the top 10 PCA of upper-triangular of connectome. In both cases, we chose as classifier a boosted ensemble of 50 shallow decision trees. For the ADNI dataset, we trained it using the RUSBoost algorithm (Seiffert et al., 2010), particularly adapted to data in which the number of input features is large with respect to the training dataset size and in which “positive” and “negative” labels are unbalanced. For the healthy aging dataset, we used a standard random forest method (Breiman, 2001). For both datasets, for training and testing we split the dataset into 5 folds, each of them with a proportion of labels maintained identical to the one of the full dataset and performed training on three of the five folds and testing on the remaining two folds (generalization performance). We considered classifiers in which the training features were of the same type of the testing features (e.g. classifiers trained on SC_emp_ and tested on SC_emp_ data; or classifiers trained on FC_MFM_ and tested on FC_MFM_ data in Figure 7D-left and 7E-right; etc.). We also considered classifiers in which the type of data differed in training and testing (e.g. classifiers trained on SC_bi-MFM_ and tested on SC_emp_ data, in Figure 7F). In all cases, generalization performance was assessed on data from different subjects than the ones used for training (i.e. prediction performed on the folds of data not actually used for training). The split in random folds was repeated 1000 times, so to be able to evaluate median performances and their confidence intervals, given by 5_th_ and 95_th_ percentile performances over the 1000 repetitions of training and testing. We measured performance based on confusion matrices between predicted and actual class labels and, just for the binary classification problem on the ADNI dataset, on the Receiver Operator Curve (ROC) analysis as well. For ROC analysis, we quantified fractions of true and false positives (numbers of true or false positives over the total number of actual positives) during generalization, which depend on an arbitrary threshold to be applied to the classifier ensemble output to decide for positivity of not of the input data. Receiver operator curves (ROC) are generated by smoothly growing this threshold. An Area Under the Curve (AUC) was then evaluated as a summary performance indicator, being significantly larger than 50% in the case of performance above chance level. The ROC curves plotted in Figure 7B and 7C, as well as their associated 95% confidence range of variation are smoothed using a cubic smoothing spline based on the cloud of TP and FP values at different thresholds over the 1000 individual training and testing classification runs. We report confidence intervals for AUCs only for “direct” classifications (pooling performances for classifiers trained on either SC_emp_ or FC_emp_ and tested on same-type empirical connectomes) and “virtual” classifications (pooling performances for classifiers trained on any type of virtual or bivirtual connectomes and tested on same nature virtual or empirical connectomes) since confidence intervals for more specific types of classifiers were largely overlapping.

**Figure 7.**
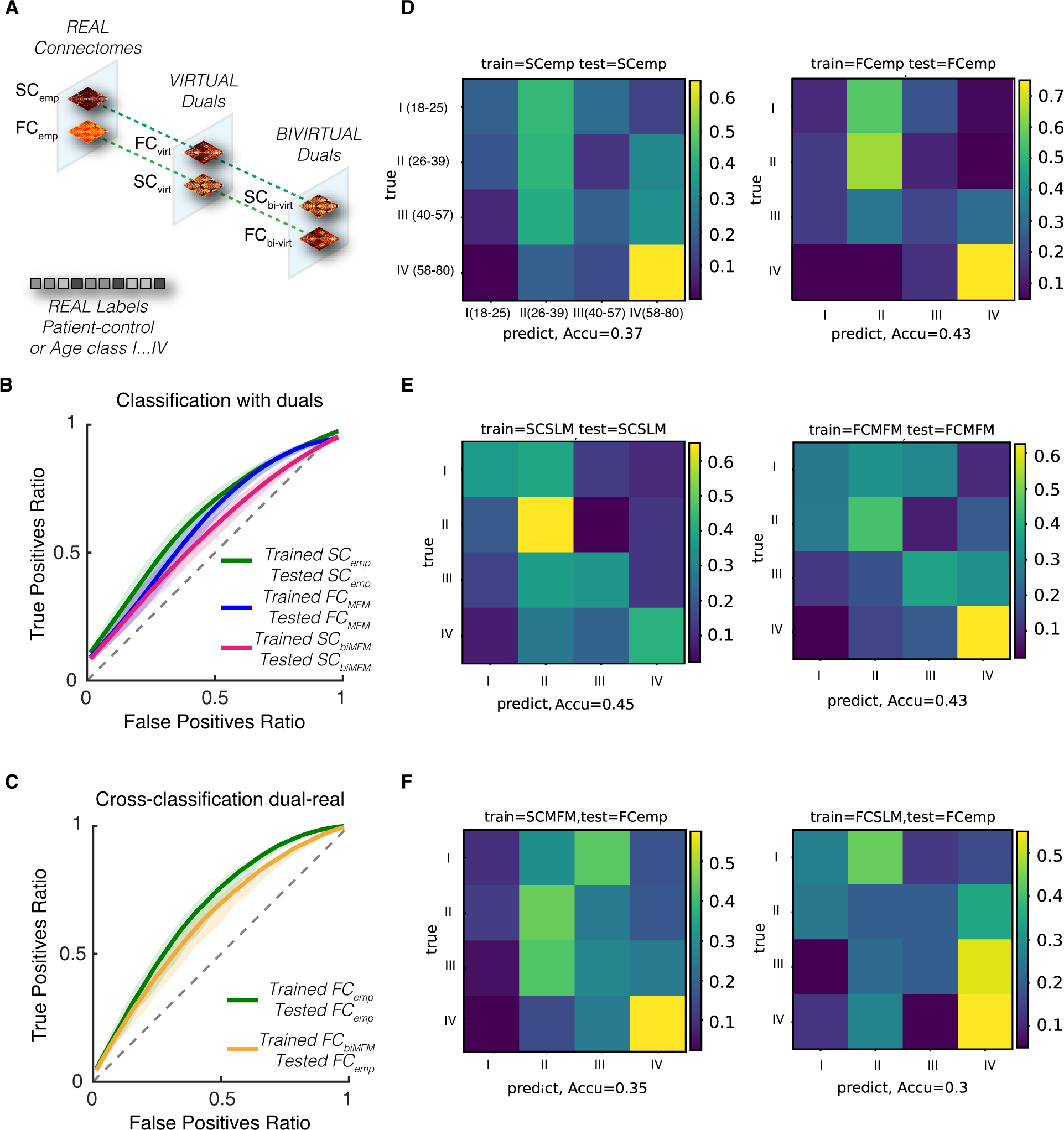
Classification of MCI patients based on empirical and virtual connectomes and virtual cohorts. A) Data completion procedures can be seen as bridges between different connectome spaces, mapping empirical connectomes in “real space” to subject-specific dual connectomes in virtual or bivirtual spaces, depending on the number of virtualization steps applied to the original connectome. Subjects classifications into controls (light blue) or MCI (yellow) and AD (red) patients are shared between empirical connectomes and their virtual and bivirtual duals. Virtual duals have a different nature than their associated empirical connectomes (empirical SCs are mapped to virtual FCs and vice versa), while bivirtual duals have the same nature. B-C) Performance of tree ensemble classifiers discriminating control from patient subjects, evaluated via Receiver Operator Curve analysis (fractions of true vs false positive, as a function of applied decision threshold; generalization performance via crossvalidation; thick lines indicate median performance, shaded regions 95% confidence intervals). In panel B, we show example of classification in dual space, compared with a real connectome space classification: in green classification with classifiers trained on empirical SCs evaluated on other empirical SCs; in blue, classifiers trained on virtual FCs evaluated on other virtual FCs (or the virtual duals of other empirical SCs); in magenta, classifiers trained on bivirtual SCs evaluated on other bivirtual SCs (or the bivirtual duals or other empirical SCs). In panel B, we show an example of cross-space classification, compared with a real connectome space classification: in green classification with classifiers trained on empirical FCs evaluated on other empirical FCs; and in orange, classification with classifiers trained on bivirtual FCs evaluated directly on other empirical FCs, without prior “lifting” into bivirtual dual space. In all the shown cases, classifications performed with classifiers trained in virtual or bivirtual connectomes are slightly less performing than for classifiers trained on empirical data, but the drop in performance is not significant for most thresholds. D-F) The confusion matrix for classification of four age classes of the healthy ageing database using the random forest Breiman algorithm is shown. D) When the classifier was trained and tested on the empirical SC and FC connectome, the accuracy was closed to ∼0.37 and ∼0.43 respectively. E) The classification accuracy for the classifier which was trained and tested on the virtual connectomes was above the chance level (∼0.25) with ∼0.43 for SC_SLM_ and ∼0.43 for FC_MFM_ connectomes which the performance was better or equivalent to the empirical connectome (D). F) Here we shown the classification performance of cross-training, when the classifier was trained on SC_MFM_ and tested on FC_emp_ with accuracy equal to ∼0.35 (F-left) and when the classifier was trained on FC_SLM_ and tested on FC_emp_ with accuracy of ∼0.30 (F-right) (see Extended Data Figure 7-1 for the classification performances on other virtual connectomes from healthy ageing dataset).

**Figure 8.**
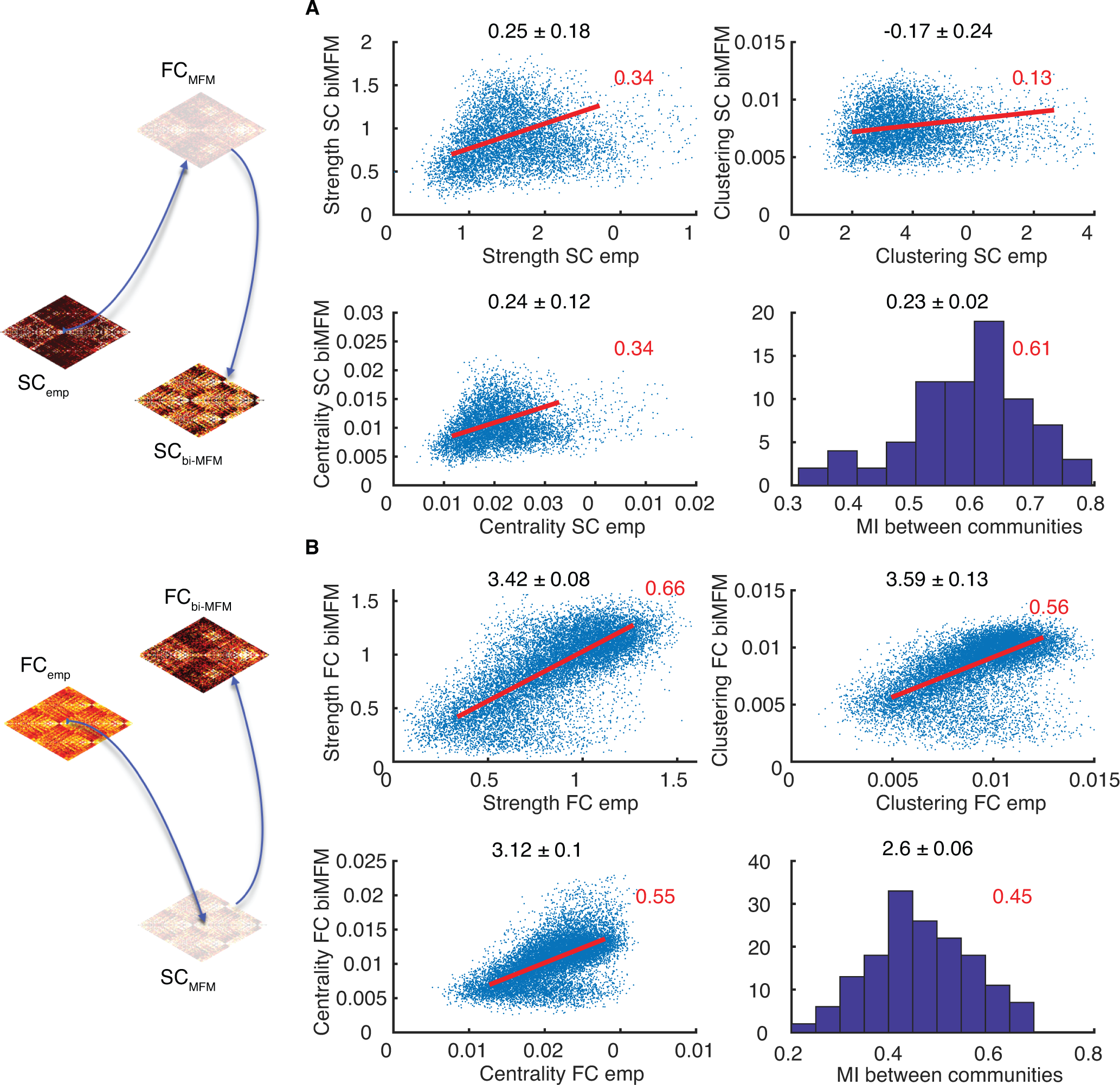
Correspondence of network topology between empirical and their bivirtual dual connectomes (ADNI dataset). The bivirtual dual connectomes share the same nature (SC or FC) of the corresponding empirical connectome. Therefore, network topology can be directly compared between empirical and bivirtual SCs or empirical and bivirtual FCs. A-B) We show here scatter plots of connectivity strengths (top left), local clustering coefficients (top right) and local centrality coefficients (bottom left) for different brain regions and subjects, plotting feature values for empirical connectomes vs their bivirtual counterparts. We also show histograms over different subjects of the relative mutual information (normalized between 0 and 1, the latter corresponding to perfect matching) between the community structures (bottom right) of empirical connectomes and their bivirtual duals. Results are shown in panel A for SC and in panel B for FC connectomes for the ADNI dataset (see Extended Data Figure 8-1 for analogous results holding for the healthy ageing dataset). In both cases, there is a remarkable correlation at the ensemble level between network topology features for empirical bivirtual connectomes (see Table 3 for the even superior correspondence at the single subject level for the ADNI dataset).

### Virtual cohorts

To generate virtual cohorts, i.e. synthetic datasets made of a multitude of virtual connectomes beyond individual subject or patient data completion, we artificially boosted the size of the original dataset by generating a much larger number of virtual subjects with multiple alternative (but all equally valuable) completions of the missing connectomic data. Concretely, to generate the virtual cohort dataset illustrated in Figure 9A, we took the 88 subjects in the SC_emp_ only plus the 12 subjects in the SC_emp_ + FC_emp_ subsets of the ADNI dataset (including 21 AD subjects, 35 MCI, and 32 Control subjects) and run for each of them the non-linear SC-to-FC completion algorithm 100 times, using each time a different random seed. The net result was a group of 100 alternative FC_MFM_ instances for each of the subjects, yielding in total a virtual cohort of 8800 FC_MFM_ matrices to be potentially used for classifier training. Such a cohort can be downloaded as a MATLAB® workspace within Extended Data FC_cohort.mat (available at the address https://github.com/FunDyn/VirtualCohorts). To generate Figure 9A, showing a dimensionally reduced representation of the relative distances between these 8800 virtual matrices, we used an exact t-SNE projection (Van Der Maaten and Hinton, 2008) of the vectors of upper-triangular parts of the different FC_MFM_ ‘s toward a two-dimensional space, using a default perplexity value of 30 and no-exaggeration.

**Figure 9.**
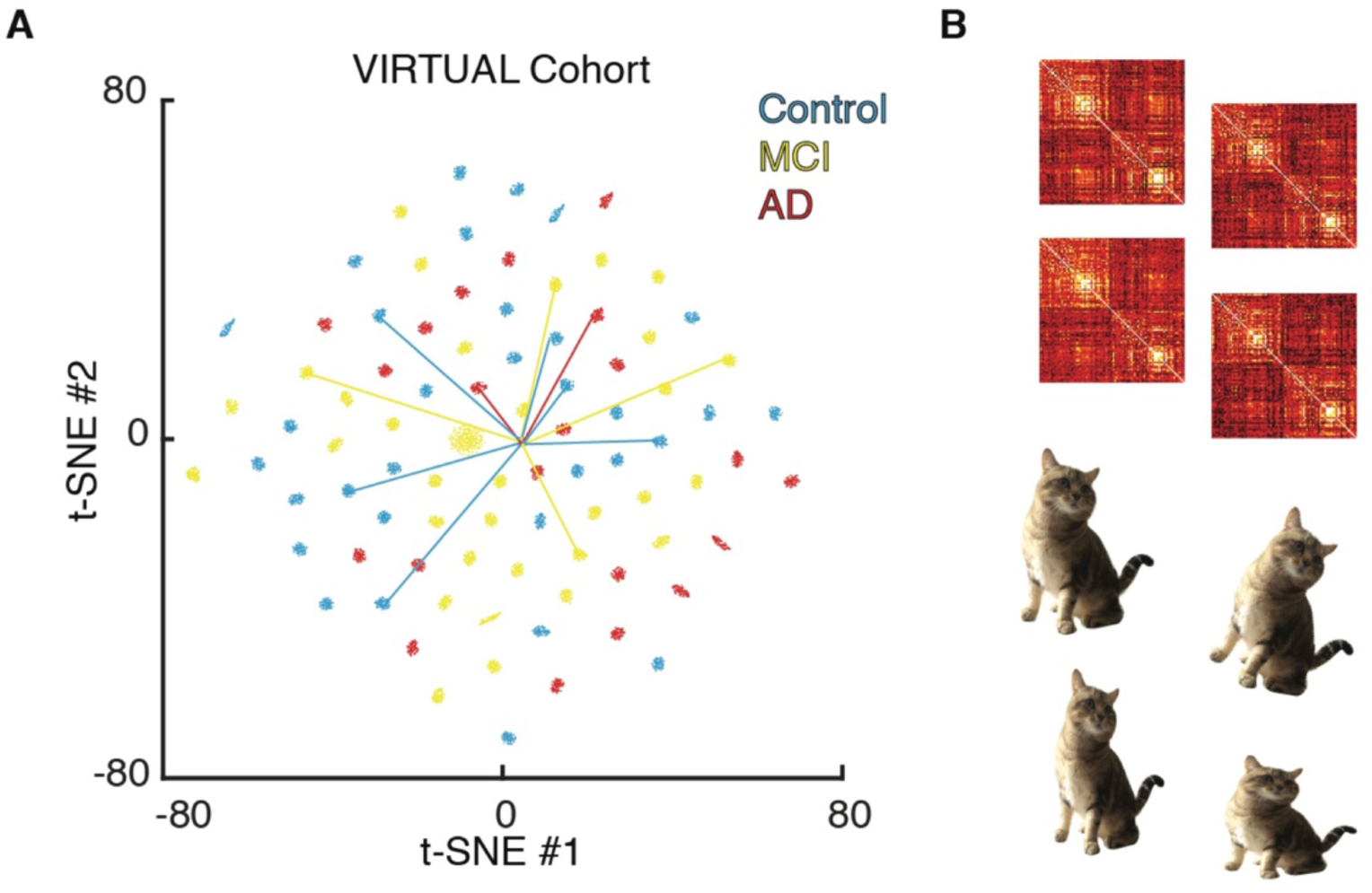
The Virtual cohorts. We created virtual cohorts of surrogate FC data, generating 100 different FC_MFM_ matrices for each of the 88 subjects in the ADNI dataset with an available SC_emp._ A) Shown here is a low-dimensional t-SNE projection of the resulting 8800 virtual FC_MFM_ ‘s, colored depending on the associated subject label (“blue” for control subjects, “yellow” for MCI patients, and “red” for AD patients). For the subjects in the ADNI “FC+SC” subset, we also projected the actual empirical FC_emp_ connectome and link their projections to one virtual connectome within the cohort for the matching subjects. All FC_emp_ connectomes appear grouped in a single cluster, since all far away to connectomes in dual space (they belong to a different space, so appear as “distant” in this projected view emphasizing differerencies within virtual space). However, virtual cohorts inter-relations reproduce an exploded view of the fine structure of this All FC_emp_ cluster. Virtual connectomes within a same virtual cohort are closer between them than connectomes belonging to different cohorts since they maintain a strict relation to their empirical counterparts and are thus good candidates for data augmentation applications. B) We show, on top, example alternative connectomes within a representative cohort for a single subject that could be used as alternative identity preserving distorted connectomes for data augmentation applications, analogously to slightly distorted versions of object images (on the bottom) used to boost training of object classifiers.

On the same t-SNE projection, beyond the FC_MFM_ connectomes within the virtual cohort connectomes we show as well additional FC connectomes, for the sake of comparison (using the same t-SNE neural network adopted for projecting the virtual cohort connectomes on the Euclidean plane). Specifically, for the 12 subjects with available FC_emp_ in addition to SC_emp_, we also show the projected positions corresponding to the real FC_emp_. Moreover, we also show positions of bivirtual FCs generated from the FC_emp_ only subset paired to the corresponding FC_emp_ projection.

## Results

### Connectomic data may have gaps: the example of ADNI

The first dataset we have chosen to focus in the framework of this study corresponds to one of the earliest and most popular available datasets in AD research, including a substantial amount of structural and functional connectomic information, i.e. the Alzheimer’s Disease Neuroimaging Initiative (ADNI) database (adni.loni.usc.edu). ADNI is impressive for the variety of features it aimed at systematically gathering (Figure 1A). Importantly, based on the T1, DTI and resting-state (rs) BOLD fMRI images available through the ADNI data-sets, state-of-the-art processing pipelines can be used to extract subject-specific Structural and resting-state Functional Connectomes, compiled into connectivity matrices adapted to the brain parcellation of choice (Figure 1B, see *Materials and Methods* for details).

We had access to 244 overall subjects (119 labeled as “MCI” and 51 as “AD”, thus 170 “Patients”, in addition to 74 control subjects, see *Materials and Methods*) for which MRI data had been gathered. We could extract an FC matrix for 168 subjects (starting from rsfMRI) and a SC matrix (starting from DTI) for 88 subjects. However, only for a minority of 12 subjects rsBOLD and DTI information were both available. In a majority of cases, either DTI or rsBOLD were missing (Figure 1C). This reduced number of “complete” subjects constitutes a serious challenge to attempts of automatedly categorize them through machine learning or inference approaches capitalizing on both SC and FC features simultaneously. As a matter of fact, the total numbers of AD- and MCI-labeled subjects in this complete subset decreased respectively to just 2 and 4, against 6 controls. In these conditions, the development of effective data completion strategies would be an important asset toward the development of classifier schemes exploiting FC/SC synergies. Therefore, approaches to “fill gaps” (completion) and, possibly, even artificially boosting sample size (augmentation) are veritably needed.

### Control dataset: healthy aging

To confirm the robustness of all following analyses performed on the first ADNI dataset, we also consider in the following comparisons with analogous analyses conducted on a second control dataset. In this previously analysed dataset (Zimmermann et al., 2016; Battaglia et al., 2020), we considered 49 healthy adult subjects covering an age-span from 18 to 80 years that we split in four age-classes (see *Material and Methods* for details). For all these 49 subjects, both FC_emp_ and SC_emp_ are simultaneously available, thus extending the number of subjects for which a ground truth connectome against which evaluate the performance of each tested completion pipeline is possible.

We also note that connectomes in the two ADNI and healthy aging datasets were defined in terms of different brain parcellations, involving a different number of regions. This fact will allow further testing the robustness of our analyses against changes of the used parcellation.

### Linking SC and resting-state FC via computational modeling

As previously mentioned, FC and SC are related only indirectly through the rich non-linear dynamics supported by brain networks (Ghosh et al., 2008; Deco et al., 2011; Kirst et al., 2016). Mean-field modeling of large-scale brain networks has emerged initially as the key tool to predict the emergent dynamic patterns of resting-state FC, from spontaneous dynamics constrained by SC (Ghosh et al., 2008). It is thus natural to propose the use of model-based solutions to perform data-completion, which, in both the SC-to-FC and FC-to-SC directions, requires to capture the inter-relation between the two as mediated by dynamics.

Large-scale mean-field brain network models are specified by: i) a parcellation of cortical and subcortical brain areas; ii) a co-registered input SC matrix in the same parcellation; iii) a forward solutions linking source and sensor space; iv) a neuronal mass model, describing the non-linear dynamics of the regions at each of the nodes of the SC matrix; v) a choice of a few global parameters (e.g. scale of strength of inter-regional connectivity or speed of signal propagation along fiber tracts); vi) an external input given to the different regions, that, in the simplest case, corresponds to simple white noise uncorrelated across each of the different sites and of homogeneous strength. The Virtual Brain enables the complete workflow from brain images to simulation (TVB; Sanz-Leon et al., 2013, 2015). Personalization is accomplished by the subject-specific structural skeleton – ingredients (i) through (iv)–, which has been demonstrated to be individually predictive (Proix et al 2017; Melozzi et al 2019). Simulations of the model can be run to generate surrogate BOLD time-series of arbitrary length (see *Materials and Methods* for details) and the associated simulated resting-state FC, time-averaged (static FC) or even time-resolved (FC dynamics or dFC, Hansen et al., 2015). The thus obtained simulated FC will depend on the chosen global parameters, setting the *dynamic working point* of the model. The model dynamics will eventually switch between alternative dynamical regimes when its global control parameters cross specific critical points. Tuning global parameters will thus uniquely determine, in which regime the model operates. Mean-field large scale models constrained by empirical SC tend to generate simulated resting-state FC that best matches empirical observations when the dynamic working point of the model lies in the proximity of a model’s critical point (Deco et al., 2011; Deco et al., 2013; Hansen et al., 2015; Triebkorn et al., 2020).

We here chose one of the simplest possible whole-brain network model designs, which emphasizes activity-based network organization (as opposed to reorganization due to synchronization) and thus ignores inter-regional propagation delays. This approach is frequently used in the literature (e.g., Deco et al., 2013; Hansen et al., 2015; Aerts et al., 2018) and has the advantage of avoiding the need for complex delay differential equation integration schemes (see *Discussion* for more details). Activation-based approaches adopt particularly simple neural mass models such as the reduced Wong-Wang model (Deco et al., 2013), in which the dynamics of an isolated brain region is approximated by either one of two possible steady states, one “down state” at low firing rate and an “up state” at high firing rate, a feature initially meant to mimic bi-stability in working memory or decision making (Wong & Wang, 2006). By varying *G* the model will switch from a low-coupling regime, in which all regional activations are low to a high-coupling regime, in which all regional activations are high, passing through an intermediate range, in which both regimes can exist in a multistable manner and regions display spatially and temporally heterogeneous activations (a changing mix of high and low firing rates). The best fit between simulated and empirical FC occurs slightly before the critical rate instability, at which modes of activity with low firing rate disappear (Deco et al., 2013).

As alternatives to the just described *non-linear mean-field models* (MFMs) of resting-state brain dynamics, simpler *stochastic linear models* (SLMs) have also been considered (Goñi et al., 2014; Messé et al., 2014; Saggio et al., 2016). In these models, the activity of each region is modeled as a stochastic process (linear, in contrast to the non-linear neural mass dynamics of conventional MFMs), biased by the fluctuations of the other regions weighted by the SC connectome (see *Materials and Methods*). SLMs have also two different regimes. In the first regime, the activities of all regions converge to a fixed-point of constant mean fluctuating activities, while, in the second, regional activities diverge with exponential growth. Once again, the best fit between the simulated and the empirical resting-state FCs is observed when tuning the model parameters slightly below the critical point (Hansen et al., 2015; Saggio et al., 2016).

MFMs and SLMs provide thus two natural ways to generate simulated resting-state FCs, depending on the chosen dynamic regime, starting from a selected SC. Strategies have also been devised to approximately solve the inverse problem of determining which SC matrix should be used as input to a model in order to give rise to a simulated FC matching a specific, pre-determined target matrix. For the SLM, a simple analytic solution to the inverse problem exists (Saggio et al., 2016). For MFMs, inverse problems have not been studied with the same level of rigor, but algorithms have been introduced that iteratively adjust the weights of the SC matrix currently embedded in the model to improve the fit between simulated and target FCs (Gilson et al., 2016; 2018). We will show later that these algorithms, although initially designed to identify changes of “effective connectivity” occurring between resting state and task conditions, have the potential to cope with the actual problem of MFM inversion, providing reasonably good ansatz for SC inference.

As linear approaches are significantly faster than non-linear approaches, it is important to study their performance alongside nonlinear approaches to confirm the actual justification of the use of more complicated algorithms. We will see that for one of the two considered datasets, the ADNI one, non-linear methods are superior for the data completion applications we are interested in. However, performance of completion happened to be slightly superior for the SLM-based than for the MFM-based methods in the case of the second healthy aging dataset (hence the interest of exploring and benchmarking both linear and nonlinear completion strategies).

### Model-driven data completion

Figure 2 summarizes many of the modeling operations described in the previous section framing them in the specific context of connectomic data completion. MRI data can be used to generate empirical SC matrices SC_emp_ (from DTI) or FC_emp_ (from rs fMRI BOLD). By embedding the empirical matrix SC_emp_ into a non-linear MFM or a linear SLM, it is possible to compute surrogate FC matrices (Figure 2A, upward arrows), denoted, respectively, FC_MFM_ and FC_SLM_. The MFM and SLM global parameters are suitably tuned (slightly subcritical) then FC_MFM_ and FC_SLM_ will be maximally similar to the empirical FC_emp_ (dynamic working point tuning, represented by dashed grey arrows in Figure 2A). Starting from the empirical matrix FC_emp_, one can then infer surrogate SC matrices (Figure 2A, downward arrows), either by using a linear theory –developed by Saggio et al. (2016)– to compute a surrogate SC_SLM_; or by exploiting non-linear effective connectivity algorithm –generalized from Gilson et al. (2016; 2018)– to infer a surrogate SC_MFM_ starting from a random initial guess (see later section).

When connectomic data are incomplete (only SC_emp_ or only FC_emp_ are available, but not both simultaneously), computational simulation or inference procedures can be used to fill these gaps: by using FC_MFM_ or FC_SLM_ as virtual replacements for a missing FC_emp_ (Figure 2B); or by using SC_MFM_ or SC_SLM_ as virtual replacements for a missing SC_emp_ (Figure 2C). The quality of the model-generated virtual SCs and FCs can be assessed by comparing them with the actual empirical counterparts for the small subset of subjects for which both SC_emp_ and FC_emp_ are simultaneously available. Optimizing the quality of the virtually completed matrices on subjects for which both empirical connectomes are available (as, e.g. the subset of ADNI “SC_emp_+FC_emp_” subjects), also allows extrapolating target criteria for identifying when the model is operating a suitable dynamic working point, that can be evaluated solely based on simulated dynamics when a fitting target matrix is missing and thus fitting quality cannot be explicitly measured (cf. Figures 3 and 4). We can thus translate these criteria into precise algorithmic procedures that inform linear or non-linear SC-to-FC and FC-to-SC completion (see Tables 1, 2 and 1-1, 2-1).

We now, provide more details on implementation and performance for each of the four mentioned types of data completion.

### Linear SC-to-FC completion

In linear SC-to-FC completion, a simple SLM (see *Materials and Methods*) is constructed based on the available SC_emp_ and its direct simulations or even, in a much faster manner, analytical formulas deriving from the model’s theory are used to generate the associated virtual Pearson correlation matrix FC_SLM_ (Extended Data Figure 3-1). In this stochastic linear modeling scheme, once the driving noise strength is arbitrarily chosen and fixed and the input connectome SC_emp_ is specified, there remains a single parameter to adjust, the global scale of long-range connectivity strength *G*. Extended Data Figure 3-1A shows a systematic exploration, performed on subjects from the ADNI “SC_emp_+FC_emp_” subset, of how the completion quality depends on tuning this parameter *G.* As shown by the main plot in Extended Data Figure 3-1A for a representative subject, increasing *G* the correlation between the empirical FC_emp_ and the simulated FC_SLM_, derived here from direct SLM simulations, initially grows to peak in proximity of a critical value *G*.* The correlation then drops dramatically when further increasing *G* beyond the critical point *G\**.

The exact value of *G\** depends on the specific personalized SC_emp_ connectome embedded into the SLM and is therefore different for each subject. The small boxplot inset in Extended Data Figure 3-1A gives the distribution of the personalized *G\** over all the subjects in the ADNI “SC_emp_+FC_emp_” subset. However, when performing linear FC completion because BOLD data and FC_emp_ are missing, the exact location of the fitting optimum cannot be determined. To perform linear SC-to-FC completion for the ADNI subjects with missing BOLD we chose to always use a common prescribed value *G*_ref_ =* 0.83, set to be equal to the median of the personalized *G\** over the “SC_emp_+FC_emp_” subset of ADNI subjects.

Once a *G*_ref_* value and a noise strength are set, the linear completion can be further sped-up by the fact that the covariance matrix FC_SLM_ for these frozen parameters can be analytically evaluated, as discussed in Saggio et al. (2016). Therefore, one can directly apply the SLM analytical formulas (see *Material and Methods*) on the available SC_emp_ as input, without the need for performing direct simulations to generate surrogate BOLD first. Extended Data Figures 3-1B-C analyze the expected performance of this “simulation-less” procedure, as benchmarked by applying it on the ADNI “SC_emp_+FC_emp_” subset. The boxplot in 3-1B (leftmost box) reports a median Pearson correlation between the linear virtual FC_SLM_ and the actual empirical FC_emp_ close to ∼0.24 for the ADNI dataset. This correlation is larger and rise to ∼0.37 for the healthy aging dataset, in which FC_SLM_ are generated from SC_emp_ using precisely the same algorithm. Panel 3-1C indicates then the percent loss in correlation that has been caused by using the common value *G*_ref_* and the analytical formula to evaluate the linear virtual FC_SLM_ rather than direct simulations at the actual personalized optimum *G\** for each of the ADNI “SC_emp_+FC_emp_” subjects. The median quality loss is approximately 0.5%, indicating that the lack of personalized tuning of the SLM working point is only a minor issue and that is acceptable to speed-up completion by relying on analytical evaluations.

Table 1-1 provides a pseudo-code for the linear SC-to-FC completion procedure (see *Materials and Methods* for all details). Linear SC-to-FC completions for the DTI-only subjects in the considered ADNI dataset and the Healthy Ageing dataset can be downloaded as part of Extended Data FC_SLM.

The median Pearson correlations of ∼0.24 or ∼0.37 between the linear virtual FC_SLM_ and the actual empirical FC_emp_ for the ADNI and the healthy aging datasets respectively are significant but still absolutely weak. A way to assess whether linear SC-to-FC completion is worthy, despite these low correlation values, it is possible to compare the achieved reconstruction quality with the one that one could trivially achieve by simply taking the SC_emp_ connectome itself as surrogate FC, since we know that SC and FC connectomes are already strongly related (Hagmann et al., 2008). This strategy of using the “other connectome” to perform FC completion would be even faster than SLM-based completion. We thus computed the percent improvement in rendering FC_emp_ via FC_SLM_ for subjects in the ADNI “SC_emp_+FC_emp_” subset and for subjects in the healthy aging datasets. As shown in Extended Data Figure 2-1A, for the ADNI dataset, the use of FC_SLM_ resulted systematically in a *worse* performance (median drop Δ_;<=>=?@_= -15%, see *Materials and Methods* for definition) in reproducing the actual FC_emp_ than using the other available connectome SC_emp_. However, in the case of the healthy aging dataset, the use of FC_SLM_ resulted in a clearly *better* performance than when using “the other connectome” (median improvement Δ_;<=>=?@_= +40%). Thus, the performance of linear SC-to-FC completion can be good but was not robustly maintained across the two considered datasets.

### Non-linear SC-to-FC completion

In non-linear SC-to-FC completion, a more complex MFM (see *Materials and Methods*) is constructed based on the available SC_emp_ and is simulated to generate surrogate BOLD data and the associated Pearson correlation matrix FC_MFM_ (Figure 3). Non-linear mechanistic MFM models are supposedly more compliant with neurophysiology than the phenomenological SLMs. Furthermore, because of their non-linearities, they are potentially able to capture complex emergent collective dynamics resulting in non-trivial dFC (which SLMs cannot render, cf. Hansen et al., 2015). However, MFMs have also more parameters and are computationally costlier to simulate than SLMs.

We chose here to limit ourselves to MFMs based on a reduced Wong-Wang regional dynamics (see *Materials and Methods* for model equations), which has previously been used to successfully reproduce rsFC (Deco et al., 2013) and dFC (Hansen et al., 2015) starting from empirical SC, despite its relative simplicity with respect to other possible neural masses implemented in the TVB platform. In addition to the global scale of long-range connectivity strength *G*, the MFM model dynamics depend also on regional dynamics parameters. In Figure 3, we froze all local parameters but the NMDA decay time-constant *τ*, since they affected the dynamic behavior of the model less than the other control parameters and, in particular, did not alter qualitatively the repertoire of accessible dynamical regimes (compare Figure 3A with Extended Data Figure 3-2). The simulated collective dynamics and the resulting non-linear virtual FC_MFM_ will depend on the choice of the free control parameters *G* and *τ*. In Figure 3A, we have explored the dependency of the correlation between FC_MFM_ and the actual empirical FC_emp_ as a function of *G* and *τ* achievable over the subjects in the ADNI “SC_emp_+FC_emp_” subset. As evident in Figure 3A, this dependence is non-monotonic and the best-fitting qualities are concentrated in a narrow concave stripe across the *G/τ* plane. Panels 3B and 3C report zoom of Figure 3A into increasingly smaller regions, revealing an extended zone of high fitting quality which some absolute optimum parameters *G\** and *τ\** (here *G\** = ∼ 1.5 and *τ\** = 25).

Remarkably, this best-fitting quality zone on the *G/τ* plane is associated as well to other properties that can be evaluated just based on the simulated dynamics (and, therefore even when the actual target FC_emp_ is unknown and missing). We found that the best fit quality systematically occurs in a region where three criteria are jointly met (Figures 3D-F).

First, there is a mixture of “ignited” regions with large activation and of not yet ignited regions with a weaker firing rate (*spatial heterogeneity*, Figure 3D). Conversely, when moving out of the best-fitting zone, the activity becomes more spatially homogeneous, either with all regions stable at low (for *G* <<< *G\**) or high (for *G* >>> *G\**) firing rates.

Second, the time-averaged FC_MFM_ has a complex modular organization between order and disorder, associated to high average clustering coefficient, in contrast with the absence of clustering observed for *G* <<< *G\** or *G* >>> *G\** (*structured FC*, Figure 3E).

Third, the simulated collective dynamics give rise to meta-stability of FC along time, i.e. to a non-trivially structured dFC, which alternates between “knots” of transiently slowed-down FC network reconfiguration and “leaps” of accelerated reconfigurations. Such non-triviality of dFC can be detected by the inspection of the so-called dFC matrix (Hansen et al., 2015; Arbabyazd et al., 2020; Battaglia et al., 2020; Lombardo et al., 2020), representing the similarity between FC matrices computed at different time-windows (see *Materials and Methods*). In this dFC matrix analysis, dFC “knots” are visualized as blocks with high inter-FC correlations, while dFC “leaps” give rise to stripes of low inter-FC correlation. The prominence of the block structure of the dFC matrix can be measured by the dFC clustering coefficient (see *Material and Methods*), higher when the dFC matrix includes more evident knots. The dFC clustering coefficient is higher in the best fit zone, while it drops moving outside it toward *G* <<< *G\** or *G* >>> *G\** (*structured dFC*, Figure 3F).

By scanning the *G/τ* plane in search of a zone with simultaneous spatial heterogeneity of activations, structured FC and structured dFC, the MFM model parameters can be tuned to bring it in a zone invariantly resulting in relatively higher fitting quality. Figure 3G shows the analysis of the expected performance of this procedure, as benchmarked by applying it on the ADNI “SC_emp_+FC_emp_” subset (on the left) and the healthy aging dataset (on the right). We measured a median Pearson correlation between the non-linear virtual FC_MFM_ and the actual empirical FC_emp_ close to ∼0.32 for both datasets, which is larger than for FC_SLM_ in the case of the ADNI but slightly maller in the case of healthy aging datasets.

Table 1 provides a compact pseudo-code for the non-linear SC-to-FC completion procedure (see *Materials and Methods* for all details). Non-linear SC-to-FC completions for the DTI-only subjects in the considered ADNI dataset can be downloaded as part of Extended Data FC_MFM.

The value of correlation with FC_emp_ achieved by FC_MFM_ can thus be larger than the one achieved by FC_SLM_ and also appear more robust, since attained in both datasets. Nevertheless, it remains necessary to check, as previously for the FC_SLM_, that it constitutes an improvement on the trivial strategy over taking the “other connectome” as substitute (i.e. taking FC to be identical to SC_emp_). In Extended Data Figure 2-1A, we show that this is indeed the case, unlike for linear SC-to-FC completion. The procedure sketched in Table 1-1 led to a median improvement on using the “other connectome” approaching ∼20% for both datasets that can go as high as +60% in some subjects.

### Linear FC-to-SC completion

In linear FC-to-SC completion, we use once again the analytic theory derived for the SLM (Saggio et al., 2016) to deterministically compute a surrogate SC_SLM_ as a function of the available FC_emp_ or, more precisely, of the resting-state BOLD_emp_ time-series used to derive FC_emp_. In this scheme, the linear virtual SC_SLM_ is indeed taken to be directly proportional to the *inverse covariance* of the BOLD time-series (see *Materials and Methods*). The proportionality constant would depend on the free parameters chosen for the SLM, serving as a link between FC and SC. Here we set arbitrarily this constant to the unit value.

Extended Data Figure 4-1 shows the analysis of the expected performance of this procedure, as benchmarked by applying it on the ADNI “SC_emp_+FC_emp_” subset. For this ADNI dataset, we measured a median Pearson correlation between the linear virtual SC_SLM_ and the actual empirical SC_emp_ close to ∼0.22. On the healthy aging dataset, this correlation rose even up to ∼0.42.

Table 2-1 provides a pseudo-code for the linear FC-to-SC completion procedure (see *Materials and Methods* for all details). Linear FC-to-SC completions for the BOLD-only subjects in the considered ADNI and the Healthy Ageing datasets can be downloaded as part of Extended Data SC_SLM.

As for SC-to-FC completions, we confirmed if the performance reached by linear FC-to-SC completion is superior to the one that is obtainable through the trivial strategy of using “the other connectome” (in this case, the available FC_emp_). In Extended Data Figure 2-1B, we show that using SC_SLM_ rather than FC_emp_ as an ersatz for SC_emp_ leads to drops of improvements in quality with a pattern similar to the reverse SC-to-FC completion, i.e. a drop in quality, with a median value of approximately -20%, for the ADNI dataset but an increase of nearly ∼50% for the healthy aging dataset. Once again, thus, linear FC-to-SC completion can yield good results, but this performance did not robustly generalize through datasets.

### Non-linear FC-to-SC completion

Non-linear FC-to-SC completion consists in the inference of a SC_MFM_ matrix that, used as input to an MFM, produces as output a simulated FC* matrix highly correlated with the available empirical FC_emp_ (Figure 4). This non-linear inverse problem is more sophisticated than linear FC-to-SC completion, because, for the MFM a theory providing an explicit formal link between input structural connectome (SC*) and output functional connectome (FC*) is not available, unlike for the SLM. Note indeed that MFMs, at the best-fitting dynamic working point, give rise not just to a single dynamical mode, but to a multiplicity of them (Deco & Jirsa 2012; Hansen et al., 2015; Golos et al., 2015) and that each of them may be associated, in general, to a different state-specific FC (Battaglia et al., 2012; Hansen et al., 2015; Kirst et al., 2016) so that the final static FC* results from averaging over a mixture of different states sampled in stochastic proportions. Therefore, to derive the FC* associated with a given input SC*, it is necessary to run explicit MFM simulations, long enough to sample a variety of possible dynamical states.

Gilson et al. (2016; 2018) have introduced iterative optimization procedures aiming at updating a current guess for the input SC* to a model in order to improve the match between the model output FC* and a target FC_emp_. They initially conceived such a procedure as a form of “effective connectivity” analysis, aiming at constructing models which capture the origin of subtle changes between resting state and task conditions. Thus, starting from an empirical SC connectivity and from a model reproducing suitably rest FC, they slightly adjusted SC weights through an iterative procedure to morph simulated FC in the direction of specific task-based FCs. Nothing however prevents to use the same algorithm in a more radical way, to grow from purely random initial conditions a suitable effective connectome, as an ersatz of missing SC_emp_, compatible with the observed FC_emp_.

In this “effective connectivity” procedure connectome weights are iteratively and selectively adjusted as a function of the difference occurring between the current FC* and the target FC_emp_. Such optimization leads to infer refined connectomes, that, with respect to empirical DTI SC matrix, may display non-symmetric connections (distinguishing thus between “feeder” and “receiver” regions as in Gilson et al., 2016) or enhanced inter-hemispheric connections, usually under-estimated by DTI (as in Gilson et al., 2018). Here we use a similar algorithm to learn a suitable non-linear virtual SC_MFM._

The initial SC*_(0)_ is taken to be a matrix with fully random entries. An MFM embedding such SC*_(0)_ is built and simulations are run to generate an output FC*_(0)_ which is compared to the target FC_emp_ of the subject for which FC-to-SC completion must be performed. The used SC*_(0)_ is then modified into a different SC*_(1)_ = SC*_(0)_ + λΔFC_(0)_ matrix, by performing a small update step in the direction of the gradient defined by the difference ΔFC_(0)_ = FC_emp_ - FC*_(0)_. A new simulation is then run to produce a new FC_(1)_. The produce is repeated generating new SC_(i)_ = SC_(i-1)_ + λΔFC_(i-1)_ until when the difference between FC_(i)_ and the target FC_emp_ becomes smaller than a specified tolerance, i.e. |ΔFC_(i)_| < ε. The last generation SC_(i)_ is then taken as non-linear virtual surrogate SC_MFM_ (see *Materials and Methods* for details).

Figure 4A provides an illustration of the nonlinear FC-to-SC completion when applied to subjects in the ADNI ADNI “SC_emp_+FC_emp_” subset. In the first step, the matrix SC*_(0)_ is random and there is no correlation between the output FC*_(0)_ and FC_emp_. Advancing through the iterations, SC*_(k)_ develops gradually more complex internal structures and correspondingly, the correlation between FC*_(k)_ and FC_emp_ increases until when it reaches the desired quality threshold, here set to CC_target_ = 0.7. This threshold quality is usually reached after ∼1500 iterations. In the ADNI “SC_emp_+FC_emp_” subset we take advantage of the availability of the actual SC_emp_ to quantify as well the convergence of SC*_(k)_ toward SC_emp_. Figure 4A shows that advancing through the iterations, the correlation between SC*_(k)_ and SC_emp_ improves, in agreement with our hypothesis that effective connectivity can provide a reasonable replacement for structural connectivity. The expected quality of reconstruction, as estimated from results on the ADNI “SC_emp_+FC_emp_” subset is reported in Figure 4B and amounts to an expected correlation between SC_MFM_ and SC_emp_ of ∼0.31. For the healthy aging dataset, we obtain a slightly smaller median value of ∼0.28, but the difference is not statistically significant.

Table 2 provides a compact pseudo-code for the non-linear FC-to-SC completion procedure (see *Materials and Methods* for all details). Non-linear FC-to-SC completions for the BOLD-only subjects in the considered ADNI dataset can be downloaded as part of Extended Data SC_MFM.

As for SC-to-FC completion, we then confirmed if the nonlinear FC-to-SC completion SC_MFM_ does provide a superior reconstruction of SC_emp_ than the trivial alternative offered by just taking the “other connectome” (the available FC_emp_). As shown in Figure 2-1B, the use of nonlinear FC-to-SC completion led to a median improvement on the order of ∼15% for the ADNI dataset and of ∼10% for the healthy aging dataset. If the improvement achieved by non-linear completion is smaller than for linear completion in the healthy aging dataset, nonlinear FC-to-SC completions succeeds in the ADNI dataset where its linear counterpart failed. Therefore, nonlinear FC-to-SC computational generation provides a worthy strategy for data completion, although not yet as efficient as SC-to-FC completion.

We note that non-linear FC-to-SC completion, as for non-linear SC-to-FC completion, is a non-deterministic procedure, meaning that a different SC_MFM_ is generated depending on the starting initial condition SC*_(0)_. However, the different non-linear virtual surrogates lie at distances from the common actual ground truth SC_emp_ which are tightly concentrated around the median correlation. As revealed by Figure 4C, the reported correlations between SC_MFM_ and SC_emp_ were within a narrow interval of ±2.5% of the relative difference from the median distance for all the tested random initial conditions (30 per subject, see *Materials and Methods*), showing that the expected performance is poorly affected by the initial conditions. This stochastic aspect of the non-linear completion algorithm is going to allow us to generate not just one but arbitrarily many completions, starting from each available empirical connectivity matrix (see later section).

### Virtual and bi-virtual duals

SLMs and MFMs have thus the capacity to bridge from SC to FC or from FC to SC in a way that, in most cases, goes beyond capturing the mere similarity between the empirical SC_emp_ and FC_emp_ connectomes. When using these models for data completion, the input matrix is always an empirical matrix (SC_emp_ or FC_emp_) and the output a surrogate virtual matrix (respectively, FC_virt_ or SC_virt_, where the index “virt” refers generally to any completion algorithm, i.e. either using the SLM or the MFM models). However, the algorithms presented in Tables 1, 2 and 1-1, 2-1 can still be applied even when the input connectivity matrix is *already* a virtual matrix. In this case, the input could be surrogate matrices (SC_virt_ or FC_virt_) from data completion and the output would be *bi-virtual* (respectively, FC_bivirt_ or SC_bivirt_), i.e. twice virtual, since, to obtain them starting from an empirical input connectome, two different model-based procedures have to be chained. The final result of passing an original empirical connectome through two chained completion procedures is then a bi-virtual surrogate matrix of the same type (structural or functional) of the initially fed connectome. In other words, SC_emp_ is mapped to a SC_bivirt_ (passing through an intermediate FC_virt_ step) and FC_emp_ is mapped to an FC_bivirt_ (passing through an intermediate SC_virt_ step). If the information loss is not too high, pairs of virtual and bivirtual SC and FC connectomes should be shared instead of pairs involving empirical connectomes, potentially reducing difficulties to disclosing in public personal clinical data (see *Discussion*).

The virtual and bivirtual matrices obtained by operations of data-completion can be seen as a set of connectomes *dual* to the original real connectome. In mathematics, one often speaks of “duality” relations when two alternative spaces are put into relation by an element-to-element structure-preserving mapping. Here, one could reinterpret our algorithmic procedures for SC-to-FC or FC-to-SC completion as mapping between alternative “spaces” in which to describe the inter-relations between the connectomes of different subjects. Although our definition of duality is not as rigorous as in more mathematical contexts (as in the case, e.g., of linear algebra dual or bidual spaces; or in graph theory, where duality refers to node-to-link transformations), we will see that dissimilarities or similarities between the personalized connectomes of different subjects are substantially preserved by the application of completion procedure that maps an original space of empirical connectomes into a dual space of virtual connectomes. In other way, the information carried by a set of connectomes and by the set of their dual counterparts is, at least in part, equivalent (cf. Figures 5, 6, 7, Table 3 and *Discussion*). In this view, the first “dualization” operation would map a real connectome to a virtual connectome of a different type (a *virtual dual*, swapping SC with FC). The second dualization would then map it to a *bivirtual dual* of the same type (mapping SC to SC and FC to FC; cf. Figure 5A-B left cartoons and 7A). If the completion quality is good, then empirical connectomes and their bi-virtual duals should be highly related between them. Before, discussing more in detail the crucial issue of the preservation or loss of personalized information in duals, we start here by performing a self-consistency check of the data completion procedures and compare thus the start (FC_emp_ or SC_emp_) and the end (FC_bivirt_ or SC_bivirt_) points of dualization chains.

Figure 5 shows the correspondence between empirical and bi-virtual SC and FC pairs, both when using SLM- and MFM-based procedures. We first evaluated the quality of SC_bivirt_ generation, over the ADNI-subset of 88 subjects for which a SC_emp_ matrix was available and over the healthy aging dataset (Figure 5A). Considering the nonlinear bi-virtual completion chain SC_emp_ to FC_MFM_ to SC_bi-MFM_ we obtained a median correlation between SC_emp_ and SC_bi-MFM_ of ∼0.58 for ADNI dataset and ∼0.64 for the healthy ageing dataset. This quality of rendering aligned well with the performance of the linear bi-virtual completion with a correlation between SC_emp_ and SC_bi-SLM_ of ∼0.63 for the ADNI dataset. On the healthy aging dataset, linear bivirtual duals SC_bi-SLM_ were of exceptionally high quality, reaching a correlation with SC_emp_ nearly as high as ∼0.92.

We then evaluated the quality of FC_bivirt_ generation over the ADNI-subset of 168 subjects for which an FC_emp_ matrix was available and over the healthy aging dataset (Figure 5B). Considering the non-linear bi-virtual completion chain FC_emp_ to SC_MFM_ to FC_bi-MFM_ the median correlation between FC_emp_ and FC_bi-MFM_ was of ∼0.59 for the ADNI dataset and of ∼0.45 for the healthy aging dataset. Moving to linear bivirtual FC_bi-SLM_, the performance on the healthy aging dataset was of ∼0.42, equivalent to the non-linear duals. However, linear bivirtual dualization failed for the ADNI dataset, with a correlation dropping to ∼0.12, not surprisingly given the poor quality of already the first step from FC_emp_ to SC_MFM._ Even in this latter case, nevertheless, the empirical-to-bi-virtual correlations remained significant.

### Are dual connectomes still personalized?

Although significant, correlations between virtual and bivirtual with matching empirical connectomes can be small. Is this average performance sufficient not to lose subject-specific information through the various steps of transformation? The most straightforward way to answer to this question is to check whether FC_(bi)virt_ or SC_(bi)virt_ connectomes are closer to the FC_emp_ or SC_emp_ of the same subject from which they derive than to the ones of other generic subjects. Since SCs and FCs are related but not identical and their divergence can be stronger or weaker depending on the subjects (Zimmermann et al., 2019) the answer to this question is not obvious and must be checked.

We therefore introduced a measure of the improvement in connectome matching obtained by using personalized virtual and bivirtual duals rather than generic connectomes. The coefficient Δ_Pers_ *(see Materials and Methods)* quantifying the percent improvement obtained by using personalized connectomes are tabulated in Table 5 for the different types of completion.

**Table 5.** Percent improvement in connectome matching obtained by using personalized virtual and bivirtual duals

| Type of completion | | $\Delta_{\text{Pers}}$ ADNI | $\Delta_{\text{Pers}}$ Healthy aging |
| --- | --- | --- | --- |
| SCemp to FCvirt | linear | $+26\% \pm 7\%$ | $+12\% \pm 4\%$ |
| | nonlinear | $+17\% \pm 5\%$ | $+13\% \pm 4\%$ |
| SCemp to SCbivirt | linear | $+40\% \pm 18\%$ | $+23\% \pm 8\%$ |
| | nonlinear | $+17\% \pm 5\%$ | $+13\% \pm 4\%$ |
| FCemp to SCvirt | linear | $+51\% \pm 35\%$ | $+28\% \pm 22\%$ |
| | nonlinear | $+200\% \pm 37\%$ | $+87\% \pm 19\%$ |
| FCemp to FCbivirt | linear | $+46\% \pm 70\%$ | $+17\% \pm 28\%$ |
| | nonlinear | $+297\% \pm 140\%$ | $+108\% \pm 52\%$ |
| FCemp test/retest $\Delta_{\text{Pers}}$ | | $+22\% \pm 13\%$ | |

*Indicated values for real/virtual and bivirtual dual are mean ± standard deviation of the mean over subjects.*

Improvements by personalization were always positive, indicating that on average some subject-specific information is preserved. These numbers, however, are diverse between datasets and completion types. Furthermore, they should be compared with the uncertainty itself existing on empirical connectomes. Indeed the Δ_Pers_ analysis implicitly assume that empirical connectomes are exact reference comparison terms. In reality, there is a strong uncertainty on empirical connectome themselves, with an elevated test-retest variability within individual subjects (Wang et al., 2012; Chen et al., 2015; Termenon et al., 2016). In particular, the connectomic dataset released together with the study by Termenon et al. (2016) allows an evaluation of what would be the expected “empirical personalization improvement” in the case in which we actually had to compare two connectomes obtained empirically for a same subject and assess how more similar are they between them, than to a connectome of the same type but obtained from a different subject. Termenon et al. (2016) considers data mediated from the Human Connectome Project and provides for 100 subjects two different FC_emp_ matrices deriving from different scans. Using a definition of the Δ_Pers_ coefficient analogous to the one used for virtual and bivirtual completions but adapted to these test-retest empirical dataset, one can estimate a value of Δ_Pers_ of about ∼+22% for empirical FCs. In other words, the similarity between two FC_emp_ from a same subject is expected to be only a 22% larger than similarity with FC_emp_ from different subjects. We do not dispose of an analogous estimation for SC_emp_ connectomes, however we expect personalization improvements to be even in this case comparable in value, if not smaller, given that inter-subject variability for SC_emp_ connectomes tend to be smaller than for FC_emp_ (Zimmermann et al., 2019).

The Δ_Pers_ registered for bivirtual dual connectomes are of the same order of magnitude than this empirical expectancy allowing us to conclude that they are “personalized” at least as much as empirical connectomes (and at least according to this rough Δ_Pers_ measure). In some cases, notably for nonlinear bivirtual FC duals, the similarity with the original empirical connectome is way larger than what expected for empirical test-retest scans, probably due to the fact, that the effective connectivity algorithm used for FC_emp_ to SC_MFM_ nonlinear completion emphasize similarities between SC and FC, thus allowing FC_bi-MFM_ to more faithfully mirror FC_emp_ without being fully identical to it (average correlation between FC_bi-MFM_ and FC_emp_ is of ∼0.4-0.6, cf. Figure 5B). Remarkably, this strong preservation of personalization by bivirtual duals is achieved despite smaller relative improvements by personalization at the first step of the dualization chain, e.g. the transition from empirical to simple virtual duals. This means that the variability generated in the simulation leading to virtual duals, although large must maintain important subject-specific features useful to regenerate a good personalization at the following stage of generating the bivirtual dual. This also means that the Δ_Pers_ measure could be a too rough and not sensitive enough metric of personalization, since it weights equally any difference or similarity in the connectomes, independently from their relevance. Better, complementary measures of personalization are thus needed.

Since individual connectomes are affected by a necessary uncertainty a more reliable measure of the quality of personalization can be achieved by looking at the capacity of dualization to preserve overall preservation of inter-subject relations rather than specific individual data-points. Indeed, individual connectomes could be distorted through the mapping into dual virtual and bivirtual spaces, but if the distortion is such to maintain the subject’s connectome close to other subjects’ connectome to which it was close and far from other subjects’s connectome from which it was far, then the possibility to discriminate subject categories based on connectome features could still be preserved. Therefore, we computed the distances between the empirical connectomes SC_emp_ (or FC_emp_) of different subjects and the inter-subject distances for corresponding pairs of subjects but, this time, between their bivirtual dual connectomes SC_bi-virt_ (or FC_bi-virt_). As shown in Figure 6 and Extended Data Table 5-1, the correlation between the inter-subject distances in real and bidual spaces were noticeable and significant, for both ADNI and healthy aging datasets and for both MFM- and SLM-based approaches (Table 5-1), apart from the very poor performance of bivirtual linear FC completion in the ADNI (expected, given previously reported failures in this case). We also noticed that distances between bivirtual duals were often amplified, with respect to the original empirical distances. The space of dual bivirtual connectomes can thus be considered as a “virtual mirror” of the real connectome space, reproducing to a reasonable extent despite some deformation of the geometry of the original distribution of subjects.

### Subject classification based on real and virtual connectomes

The compilation of large datasets, including connectivity data from structural and functional neuroimaging is considered essential for the development of algorithmic patient stratification and predictive approaches. Here, we have described approaches for connectomic data completion and studied their consistency. We now show that such completion procedures are also compliant, in perspective, with the extraction via machine learning algorithms of the personalized information preserved in duals.

As a first proof-of-concept, we studied here two simple (and academic) supervised classification problems in which subjects are separated into different classes based on connectomic features –empirical and/or virtual– used as input. First, in the ADNI dataset, we try separating subjects into two subgroups of *control* and *patients* (i.e., MCI *or* AD) subjects. Second, in the healthy aging dataset, we separate subjects into four classes of age, from the youngest to the oldest. Importantly, input features can be computed from all different types of connectomes: (at least for the subjects for which they were available): empirical SC_emp_ or FC_emp_; their virtual duals FC_MFM_ or SC_MFM_; or their bivirtual duals SC_bi-MFM_ or FC_bi-MFM_ (see Figure 7).

### Discriminating control and patient subjects in the ADNI dataset

For the first toy classification problem, we used target classification labels already provided within the ADNI dataset, assuming them to be exact (see *Materials and Methods* for a summary of the used stratification criteria). We performed then classification based on input vectors of regional *node strengths* estimated subject-by-subject from the connectome matrices of interest (*Q =* 96 input features, corresponding to the number of brain regions in the used parcellation, see *Materials and Methods*). As supervised classifier algorithm, we chose a variant (Seiffert et al., 2010) of the random forest algorithm, which is particularly suitable when the number of input features is of the same order of the number of available data-points in the training set (Breiman, 2001), as in our case.

Examples of ADNI classifications based on empirical connectomes are shown in Figures 7, notably, based on SC_emp_ matrices (green line, Figure 7B) or on FC_emp_ matrices (green line, Figure 7C). The available subjects were randomly split into a training set and a testing set (with maintained relative proportions of the different classification labels). Figures 7B and 7C describe the average generalization performance for classifiers trained on the training set and evaluated on a testing set. Training and testing on real empirical connectomes, we achieved a moderate but significantly above chance level classification performance, as revealed by the green Receiver-Operator-Curves (ROC) in Figures 7B and 7C, for both SC_emp_ and FC_emp_ connectomes, deviating away from the diagonal (corresponding to chance level classification performance). As a more quantitative measure, one can also measure the median Area Under the ROC Curve (AUC), here equal to ∼0.69 for the SC_emp_ on SC_emp_ classifier and to ∼0.75 for the FC_emp_ on FC_emp_ classifier. AUC scores for different types of classification on the ADNI dataset are compiled in Extended Data Tables 3-1 and 3-2.

We considered then ADNI classification based on virtual and bivirtual duals instead of empirical connectomes. In this case of “dual space classification” (Figure 7B), virtual and bivirtual duals are used both when training the classifiers and when evaluating them. Therefore, to classify a new empirical connectome with a “dual space classifier”, it is first necessary to “lift” it in dual space, i.e. to map it via data completion algorithms to the suitable type of dual for which the classifier has been trained. Figure 7B shows two examples of dual space ADNI classification based on FC_MFM_ (blue curve, median AUC ∼0.64) and SC_biMFM_ (magenta curve, median AUC

∼0.59), respectively virtual dual and bivirtual duals of the real connectomes SC_emp_. Once again, for both virtual and bivirtual duals, classification performance remained above chance level. While the classification performance drops slightly with respect to classification with the actual empirical connectomes, this drop was not significant for a broad range of the most conservative decision thresholds. Above chance-level classification is thus possible as well using dual connectomes generated from data completion, achieving performances substantially equivalent to the one obtained for empirical connectomes.

We considered finally the case of ADNI classifiers trained on bivirtual duals and then evaluated on empirical connectomes (Figure 7C). In this case of “cross-space classification”, the trained classifier is able to operate in a performing manner as well on a different type of connectomes (e.g. empirical) than the one for which it has been trained (e.g. bivirtual dual). Therefore, to classify a new empirical connectome with a “cross-space classifier”, it is not necessary to first lift in dual space as for dual space classifiers. Figure 7C shows an example of cross-space classification trained on bivirtual dual FC_biMFM_ and then tested on FC_emp_ (orange curve, median AUC ∼0.70). Remarkably, the performance was not significantly different for most decision thresholds from classification trained and tested on empirical FC_emp_ connectomes. Therefore, classification of empirical connectomes based on classifier trained on virtual connectomes is possible as well.

Significant classification was possible even for some other combinations of connectomes (see Extended Data Tables 3-1 and 3-2), however performance was poorer in most cases. We did not attempt classification based on SLM-based virtual and bivirtual duals, given the deceiving quality of connectome rendering by these linear methods (in the ADNI dataset).

### Discriminating age classes in the healthy aging dataset

For the second toy classification problem, we split the subjects in the healthy aging datasets into four age categories and used the ordinal number of the age class from I to IV as target classification label. As input features we did not use any more high-dimensional vectors of connection strengths but the loadings on the first 10 principal components of each connectivity matrices. As classifier we still used random forests Breiman algorithm (see *Materials and Methods* for full detail). As before, we highlight here a few examples of classification with real empirical connectomes (Figure 7D), classification in dual space (Figure 7E) and cross-space classification (Figure 7F). We characterize performance both in terms of general accuracy (fraction of subjects correctly classified in their age class) and of detailed confusion matrices between the actual and the predicted age classes, revealing typical error syndromes. General accuracies were typically above the chance level of ∼25%, approaching (or exceeding), for instance, ∼37% for classifiers: trained and tested on SC_emp_ (Figure 7D, left, ∼37% accuracy) or FC_emp_ (Figure 7D, right, ∼43% accuracy); or, in virtual dual space, on SC_SLM_ (Figure 7E, left, ∼45% accuracy) or FC_MFM_ (Figure 6D, left, ∼43% accuracy). For cross-space classification examples, accuracies dropped but remained, e.g., of ∼35% for classifiers trained on SC_MFM_ and generalized on FC_emp_ (Figure 7F, left) or of ∼30% when trained on FC_SLM_ and tested on FC_emp_. More examples are shown in Figure 7-1, including for classifiers using bivirtual connectomes (e.g. classifiers trained and tested on FC_bi-SLM_ with an accuracy of ∼42%; but a minority of classifications were below chance level, e.g. trained on FC_bi-SLM_ and tested on FC_emp_, with an accuracy of only ∼19%).

General accuracy does not reflect fully the performance, since it averages over all possible classes. The capability to proper classify subjects of specific classes could be much larger. For instance, all but one of the classifiers highlighted in Figures 7D-F would classify elderly subjects in the IVth age class (58-80 yrs) with accuracies exceeding ∼60%. Furthermore, when misclassified, subjects tended to be attributed to neighboring but not radically different age classes –e.g. class I (18-25 yrs) with class II (26-39), or class IV (58-80 yrs) with class III (40-57)–, more rarely mixing up classes with stronger age separation. Such misclassification may also reflect meaningfully differences between subjects, whose connectome could look “younger” or “older” than the median of their age class, possibly reflecting cognitive differences, large within each age class (cf. Glisky, 2007; Battaglia et al., 2020). The analysis of factors explaining misclassification goes however beyond the scope of the present study.

As a matter of fact, we are still far from providing authentically useful examples of classification, neither on the ADNI dataset nor on the healthy aging dataset. However, this was not our aim here, the chosen classification problems themselves being rather academic and serving as first proofs-of-concept. Importantly, we can at least show that dual and cross-space classification performance, if not good, was not much worse than for real empirical connectomes. This step is already sufficient to show that empirical and virtual duals share an extractable part of information and that this shared information can be still relevant for classification.

Such information preservation, despite loose correspondence, can be explained by revealing the similarity of network topology features between real connectomes and their bivirtual duals, independently from our capacity to achieve more or less performing classifications based on these features.

### Matching network topology between real and virtual connectomes

The connectome matrices describe the weighted undirected topology of graphs of structural or functional connectivity. All information conveyed by these connectomes about pathology or other conditions is potentially encoded into this network topology. While genuine model-free analyses of network topology across all scales are still under development –see for instance, promising topological data analyses approaches (Petri et al., 2014; Sizemore et al., 2018)–, classic graph theoretical features provide a first multi-faceted characterization of the specific features of each individual connectome object (Bullmore & Sporns, 2009). We evaluated here for each empirical connectome SC_emp_ or FC_emp_ a spectrum of different graph theoretical features. In particular we evaluated for both the ADNI and the healthy aging datasets and for each brain region within each of the connectomes (see *Materials and Methods* for details): the total *strengths* (sum of the connection weights of all the links incident the region); the *clustering coefficients* (tendency of the regions neighboring to the considered node to also be interconnected between them); and *the centrality coefficients* (tendency for any path linking two different nodes in the network to pass through the considered node), evaluated via the PageRank algorithm (Brin & Page, 1998). We also evaluated for each connectome its *modular partition* into communities, by using a Louvain algorithm with default parameters (Blondel et al., 2008). Finally, we also inspected the global link *weight distributions.* We then evaluated analogous quantities for the dual connectomes associated with each of the connectomes, focusing here, for conciseness and simplicity, on bivirtual duals, sharing a common nature (Structural or Functional) with their correspondent empirical partner.

In Figure 8 we illustrate this correspondence between graph-theoretical features evaluated for different real/bivirtual dual connectome pairs in the ADNI dataset. An analogous figure for the healthy aging dataset is shown in Figure 8-1, showing qualitatively equivalent results. To compare node degrees, clustering and centrality features we plot, for every brain region in every connectome, the feature value evaluated in a real connectome against the corresponding feature value evaluated in the associated bivirtual dual. To compare community structures, we evaluate for every real/bivirtual dual connectome pair the relative mutual information MI normalized by entropy H (see *Materials and* Methods) between the community labels extracted for the two connectomes, with 0% ≤ MI/H ≤ 100% and 100% corresponding to perfect overlap. We show results for ADNI (or healthy aging) SC real/bivirtual dual pairs in Figure 8A (Figure 8-1A) and for FC pairs in Figure 8B (Figure 8-1B). In all cases we find correspondence between real and bivirtual dual connectome features significantly above chance levels. Highly significant real/bivirtual dual correlations subsist for regional strengths and centralities. For ADNI FC, these correlations can become as high as CC_median_ = 0.66 (95% bootstrap confidence interval) for regional strengths and CC_median_ = 0.55 (95% bootstrap confidence interval) for regional centralities. Correlations are found even for regional clustering coefficients, even if the small values of clustering coefficients observed in SC_emp_ connectomes are systematically overestimated in the denser bivirtual dual SC_biMFM_. Finally, concerning community matching, for SC and FC real/bivirtual dual pairs we found a median relative mutual information of ∼61% and ∼45% respectively, for the ADNI dataset, safely above chance level (estimated at ∼16%, permutation-based 95% confidence interval). (see Table 3 for the superior correspondence at the single subject level). For the healthy ageing dataset, for both SC and FC these correlations were even higher (Figure 8-1) with CC_median_ ≈ 0.8 for regional strengths, centralities, and clustering coefficients of SC real/bivirtual dual parts and CC_median_ ≈ 0.7 for the FC real/bivirtual dual parts. Finally, for the community matching for SC pairs the median relative mutual information was ∼44% and for FC pairs ∼50% (see Table 4 for the superior correspondence at the single subject level for healthy ageing dataset).

The analyses of Figure 8, and Figure 8-1 are performed at the ensemble level, i.e. pooling network features estimated from different subjects into a same point cloud. However, network features can have important variations of values not only across regions but also across subjects, which is expected to be a key indicator of subject-specific traits useful for classification. The capability to preserve these traits would thus be a crucial factor allowing the achievement of personalization when generating virtual and bivirtual duals. Therefore, we computed correlations between vectors of regional features in real and empirical connectomes but now limited to be *within* individual subjects obtaining thus, for every feature type, a different correlation value for every subject. Table 3 (for the ADNI dataset) and Table 4 (for the healthy aging dataset) show that within-subject correlations were also high (apart for SC clustering) and, for FC, even superior to ensemble-level correlations, manifesting, once again, the personalized nature of bivirtual dual connectomes. Indeed, when computing personalized correlations for pairs of real and bivirtual connectomes associated to a same matching subject, they resulted systematically superior to unpersonalized control correlations evaluated over real/bivirtual connectome pairs assembled out of different subjects (see *Materials and Methods*). Percent improvements in same-subject real/dual correlations with respect to average correlations in cross-subject pairs are compiled as well in Table 3 and Table 4. Personalization can lead to very strong percent improvements in real/virtual topology correlations, particularly in the case of FC connectomes. The operation of dualization thus preserves aspects of network topology which are specific to each subject and not just generic to a connectome ensemble.

Finally, we plot in Figure 6-1, global distributions of link weights for the different types of connectomes and both datasets. Most distributions displayed an overall similarity in shape: SC weights distributions with a peak at small values and a fat right tail; FC weights distribution more symmetric and with a broader peak at intermediate strengths. These different distribution shapes reflect that SC_emp_ networks are diluted matrices with a few strong connections only, while FC_emp_ networks have a higher and more uniform density of connections. Virtual and bivirtual SC connectomes tend to have fatter right tails (and even displaced mode peaks for SC_MFM_), reflecting that, in absence of any arbitrary sparsification strategy, completion pipelines generate surrogate SCs without the sparsity constraint and, thus, with less near-zero link weights. Such systematic discrepancy, well visible in Figure 6-1, however, does not prevent correlations between single subject-specific connectivity traits to remain strong, which is a necessary condition for personalized predictive information preservation.

### Virtual cohorts

All nonlinear data completion algorithms involve a stochastic component. Therefore, by construction, each simulation run will provide different virtual and bi-virtual connectomes, associated with the same empirical seed connectome. This property allows the generation of an arbitrarily large ensemble of surrogate virtual connectomes, forming the *virtual cohorts* associated with a specific subject (see *Materials and Methods*). Every virtual cohort maintains a strict relation to its empirical counterparts because all the matrices in the cohort are dual to the same original empirical connectome. In particular, distances between virtual connectomes sampled within two different virtual cohorts were always closely correlated to the distance between the respective seed connectomes of the two cohorts. The close relationship between the original data and the respective virtual cohorts (already studied in Figure 6 for individual instances of bivirtual connectomes) is visually manifested in Figure 9A where a distance-respecting non-linear t-SNE projection (Van Der Maaten & Hinton, 2008) has been used to represent in two dimensions the virtual cohorts of surrogate virtual FC_MFM_’s associated to the 88 subjects with available SC_emp_ in the ADNI dataset (among which, thus, also the 12 of the “SC+FC” subset). Every dot corresponds here to the two-dimensional projection of a high-dimensional virtual dual FC_MFM_ (100 different virtual FC_MFM_’s have been generated starting from each one of the 88 SC_emp_ connectomes). Clusters of dots (color-coded by their nature, of control subjects or MCI and AD patients) are visually evident in the projection indicating that the distance between dual connectomes within each virtual cohort is smaller than the distance between dual connectomes belonging to different cohorts.

We also plotted, for comparison, the cloud of the projected FC_emp_ connectomes for the twelve subjects of the ADNI “SC+FC” dataset for which it was available, and connected these projections via a thin line to the projection of one of their virtual FC_MFM_ images in the corresponding subjects’ virtual cohorts. The projections for all the FC_emp_ connectomes seem to collapse in a single additional cluster close to the center of the global t-SNE map. This collapse manifests that empirical connectomes and virtual connectomes live in different spaces, as previously stressed (Figure 7A). Eventually, when projecting a sample composed of hundred more virtual than empirical connectomes, the two-dimensional rendering of the original high-dimensional metric relations is dominated by virtual connectomes. Therefore, the cloud of the empirical connectomes’ projections appears, using a figurative image, as a “distant galaxy”, with the dots (“stars”) associated to different subjects appearing grouped in a small region of the observation field. Nevertheless, the distances between stars within the distant galaxy are mirrored by the distances between the foreground FC_MFM_ cohorts “globular clusters” mapped to each of these distant background FC_emp_ stars. The thin lines linking FC_emp_ to one of their FC_MFM_ images reveal indeed the global t-SNE projection contains an exploded view of the projection of the original “SC+FC” subset FC_emp_ connectomes (further confirming for virtual cohorts the preservation of inter-subject distances in bivirtual duals revealed by Tables 3 and 4).

A further analogy could be drawn between generating a cohort of virtual connectomes rather than a single virtual connectome and between generating an ensemble of slightly rotated or distorted images (Figure 9B). Different connectomes in a same cohort could be conceptualized as different “views” of the same connectome (as the four representative connectomes in the top of Figure 9B, sampled within the cohort of a same subject) much like different transformations of a single image that modify the exact appearance but do not prevent losing the identity of the depicted object (as the four warped kittens at the bottom of Figure 9B). For these reasons, the generation of virtual cohorts including a larger number of identity-preserving redundant connectome items may become in perspective beneficial to classifiers training, as a form of “data augmentation”, commonly used in machine learning applications in image recognition (Taylor & Nitshcke, 2018; see *Discussion*).

## Discussion

We have here demonstrated the feasibility of connectomic dataset completion using algorithms based on mean-field computational modeling. In particular, we have completed an ADNI gold standard connectomic dataset and verified that analogous completion performance could be reached on a control healthy aging dataset. We have then shown that machine learning classifiers trained on virtual connectomes can reach comparable performance to those trained on empirical connectomes. This renders the classification of novel empirical connectomes via classifiers trained exclusively on virtual connectomes possible. Furthermore, the generation of virtual and bivirtual dual connectomes is a procedure preserving at least some personalized information about detailed network topology. As a consequence, virtual cohorts offer an immense opportunity to enable or unblock, and, in perspective, possibly improve machine learning efforts on large patient databases.

Incomplete datasets for clinical research are certainly among the factors contributing to slow progress in the development of new diagnostic and therapeutic tools in neurodegenerative diseases and Alzheimer’s disease (AD) in particular. Our data completion procedures provide a step forward toward “filling dataset gaps” since they allowed us to infer Functional Connectivity when only Structural Connectivity was available or Structural Connectivity (SC) when only Functional Connectivity (FC) were available. Such procedures for data completion could easily be implemented within popular neuroinformatic platforms as The Virtual Brain (TVB). TVB provides practical graphical interfaces or fully scriptable code-line environments for “plug-and-play” large-scale brain network behavior, signal emulation, and dataset management, including simulating SC and FC with adjustable complexity MFMs or SLMs (Sanz-Leon et al., 2013). In this way, capitalizing on the software built-in capabilities, even the more elaborated non-linear completion algorithms could become accessible to non-expert users with only a little training. The possibility of having access to both types of connectomic information brought up by model-based data completion is vital because structural and functional connectivity convey complementary information. It has been shown for instance, that analyses of SC-to-FC inter-relations can yield better characterizations and group discriminations than analyses of SC or FC alone in a variety of pathologies or conditions (Zhang et al., 2011; Davis et al., 2012; Zimmermann et al., 2016; Straathof et al., 2019).

Indeed, FC networks in the resting-state do not merely mirror SC but are believed to be the by-product of complex dynamics of multi-scale brain circuits (Honey et al., 2007; Deco et al., 2011). As such, they are constrained but not entirely determined by the underlying anatomy (encoded in the SC matrix), as also confirmed by the fact that variability between FCs of different subjects may be larger than the one between SCs (Zimmermann et al., 2019). Indeed, FC also carries valuable information about the dynamic regime giving rise to the observed resting-state activity fluctuations (Hansen et al., 2015) and FC differences are thus leveraged by the nonlinear effects of dynamics that small variations in SC can have and that MFM models can in principle capture.

In particular, brain networks are thought to operate at a regime close to criticality. For a fixed SC, the resulting FC would be different depending on how closely dynamics is tuned to be in proximity of a critical working point (Deco et al., 2013; Hansen et al., 2015). This information that brain networks are supposed to operate close to a critical boundary is used to generate the surrogate virtual FC_MFM_, when performing non-linear SC-to-FC completion. Thus, FC_MFM_ carries indirectly extra information about a (putative) dynamic regime that was not conveyed by the original empirical SC (nor by virtual completions with linear SLM-based pipelines). This effective “reinjection” of information could potentially compensate for unavoidable loss –cf. “data processing inequality” (Cover & Thomas, 2006)– along the algorithmic processing chain represented by completion. This could be a possible explanation for the superior performance of nonlinear methods in the ADNI dataset completion. For this compensation to happen, however, the guess about the right working point should be close to reality. In this paper we were implicitly supposing that all the subjects have the same working point of dynamic operation (e.g. the same distance from critical rate instability, Hansen et al., 2020). Now, pathology or aging may precisely be also altering this working point itself, making of our assumption in MFM-based completion only an approximation. For instance, the distributions of matching between empirical and virtual community structure in FC connectomes for the healthy aging dataset (Figure 8-1B) are clearly bimodal, indicating that the used completion ansatz may be more appropriate for certain subjects than for others. Thus, diverse working points of dynamic operation for different subjects, here not accounted for, may contribute to the inferior performance of nonlinear methods in the healthy aging dataset. We defer to future studies considerations about how to further optimize the selection of a working point.

When both empirical SC and FC were available, we could measure the quality of reconstruction achieved by our models. The correlation reached between empirical and reconstructed connectivity matrices is only moderate, however. There are multiple reasons for this limited performance. One evident reason is the simplicity of the neural mass model adopted in our proof-of-concept illustration. The Wong-Wang neural mass model is able only to express two states of lower or higher local activation (Wong & Wang, 2006). Instead, neuronal populations can display a much more extensive repertoire of possible dynamics, including e.g., coherent oscillations at multiple frequencies, bursting, or chaotic trajectories (Stefanescu & Jirsa, 2008; Spiegler et al., 2011). Synchronization in a network depends on various factors, including frequency, network topology, and time delays via signal propagation, all of which have been ignored here and in large parts of the literature (Deco et al., 2009; Petkoski & Jirsa, 2019). It is acknowledged that delay-less approaches serve as a useful approximation (Deco et al. 2015). Nevertheless, we are aware that our choice to restrict our analyses on the subset of activation-based mechanisms introduces critical limitations. Indeed, our models, ignoring delay-mediated synchronization, are incapable of capturing a range of dynamic oscillatory behaviors, such as multifrequency coupling or multiphase coupling. More sophisticated mean-field virtual brain models could thus reach superior performance (see e.g. Stefanovski et al., 2019), going beyond the first proof-of-concept examples presented here.

Yet, even such a simple model, achieving such a limited reconstruction performance proved to be consistent and useful. First, when concatenating data completion pipelines to give rise to bi-virtual data, we found a robust self-consistency, i.e. remarkable matching between e.g. the original SC (or FC) and the bi-virtual SC_bi-MFM_ (or FC_bi-MFM)_ generated via the intermediated FC_MFM_ (or SC_MFM_) step. This self-consistent correspondence is not limited to generic correlations but captures actual personalized aspects of detailed network topology (Table 3 and Figure 8 for the ADNI dataset and Table 4 and Extended Data Figure 8-1 for the healthy ageing dataset). Second, classification performance reached based on empirical data could be nearly equated by classifiers trained on virtual or bivirtual dual connectomes (Figure 7). Therefore, even if the reconstruction quality of our model-based completion procedures is modest, a meaningful relationship with the original seed data is still maintained, even after two steps of virtual completion. The use of simple models has the additional advantage of being less computationally expensive to simulate. SLMs are even simpler and faster to run than our basic MFMs and their performance was better than the one of nonlinear models in many aspects when dealing with the healthy aging dataset. Note that SLMs have been shown to be very performing in rendering static aspects of FC in other contexts as well (Hansen et al., 2020; Messé et al., 2014). However, linear models were down-performing on the ADNI dataset, while nonlinear models performance seemed more stable across datasets. This shows once again that linear and nonlinear models may capture different facets of the actual, possibly unknown empirical connectomes and that there is an interest in computing and sharing both type of surrogates, given their potential complementarity.

In terms of computation costs, basic MFMs as our virtual brains based on the Wong-Wang model, provide a reasonable compromise between computational speed and the need to render structured brain dynamics beyond mere Gaussian fluctuations (Haken, 1983) constrained by SC. The most expensive aspect of nonlinear completion procedures –both SC-to-FC and FC-to-SC– is however their iterative nature. Indeed, not just one, but many virtual brain simulations must be performed, to scan parameter space for the best working point for FC simulation (cf. Figure 3) or to grow from random initial conditions an effective connectivity matrix sufficiently mature to render genuine aspects of SC (cf. Figure 4). Note however that, in reality, the number of iterations can be dramatically reduced by choosing good guesses for initial conditions. In the case of SC-to-FC completion, the a priori knowledge that best working point lie close to a critical line and that the monitored metrics landscape is convex, a bisection search strategy (Boyd & Vanderberghe, 2004) can be used instead of exhaustive grid search. In the case of FC-to-SC completion, starting from an initial SC* conditions close to a generic group-averaged SC connectome rather than fully random can speed-up convergence.

We have provided in Figure 7 the first proof of concept of the possibility to use virtual and bivirtual connectomes for performing subject classification. For the purpose of classification, data completion procedures are seen as veritable computational bridges between alternative “spaces” in which to perform machine learning, linked by duality relations (Figure 7A). We propose in this respect two possible types of strategy. The first one is to abandon the “real space” of actual empirical connectomes and to operate directly in dual spaces (Figure 7B). In these approaches, empirical connectomes would have to be transformed into their virtual or bivirtual dual counterparts as a necessary pre-processing step. In the second type of strategy, classifiers trained in dual spaces are used to operate in the real space. While such approach doesn’t require the virtualization of empirical input connectomes prior to their classification, performance could be potentially reduced by a possible systematic mismatch in input feature distributions between real and dual spaces (Figure 8 and Extended Data Figure 8-1 show, for instance, some network features such as, respectively, SC clustering or SC weights themselves tend to get overestimated in dual connectomes). The specific examples highlighted in Figures 7B and 7C for ADNI patient discrimination and Figures 7D-F for healthy aging age class prediction show comparable qualities of classification for dual space and cross-space classifications (in both cases, not significantly decreases with respect to classification in real space). Generally, we were able only to reach poor classification performances, barely above chance level. However, the performance was not significantly better for direct classification based on empirical connectomes. As a matter of fact, we have to acknowledge that we are still far from being able to reliably discriminate subject classes based on connectome features, independently from training being performed on real or dual connectomes. We would like to stress that the number of used input features –e.g. *K =* 96, corresponding to the number of regions in the used parcellation (see *Materials and Methods*) for which connectivity strengths were computed in the ADNI dataset classification problem – is comparable to the number of subjects in the considered dataset (*N =* 88 or 178 respectively for ADNI subjects with available SC_emp_ or FC_emp_). Therefore, it is not surprising that high performances are difficult to access, even when using classification approaches specially adapted to this situation, as in our case. Superior classification performance could be potentially reached via a more careful feature selection (Guyon & Elisseeff, 2003) that goes beyond the scope of the current study. Hopefully, future attempts to classification will be able to approach more robustly these tendential performances. Given the high degree of personalized correspondence between real and dual connectomes (cf. Table 3 for the ADNI dataset and Table 4 for the healthy ageing dataset), we are confident that any performance level reached by future classifiers trained in real space could be closely approached by classifiers trained in dual virtual and bivirtual spaces.

In perspective, the use of virtual connectomes could become beneficial to the training of machine learning algorithms in a further way. The use of a wider ensemble of surrogate date with statistical distributions of multi-dimensional features equivalent to the original data is a common practice in machine learning, known as *data augmentation* (Yaeger et al., 1997; Taylor & Nitshcke, 2018), as previously mentioned. Data augmentation is e.g. very popular in object recognition (where surrogate training data are produced by clipping or variously transforming copies of the original training images). Data augmentation aims to expand the training dataset beyond the initially available data to boost the learning by a classifier of the target categories (e.g. object identities). Crucial for dataset augmentation applications is that the surrogate data generated are not just identical to the actual data with some added noise but are genuinely new and can serve as actual good guesses for alternative (unobserved) instances of data-points belonging to the same category (cf. Figure 9B). Indeed, if information cannot be created (Cover & Thomas 2006), redundant information can nevertheless improve the performance of decoding and classification (Guyon & Elisseeff, 2003). Computational models such as MFM do not provide mappings between input and output connectomes, but rather between statistical ensembles of connectomes, with both mean and correlated dispersion realistically shaped by trustworthy non-linear dynamics. In other words, differences between alternative connectomes in a generated surrogate virtual cohort are not mere “noise”, but reflect realistic data-compliant possibilities of variation. The different connectome realizations sample indeed the specific landscapes of possible FCs that may be compatible with a given SCs, degenerate because the allowed dynamics to unfold along with low-dimensional manifolds, rather than being frozen in strict vicinity of a trivial fixed point (Mehrkanoon et al., 2014; Pillai & Jirsa, 2017). Therefore, given that inter-relations between virtual cohorts mirror inter-relations between empirical subjects (Figures 6 and 8, Extended Data Figure 8-1, Tables 3, 4, 5, and Extended Data Table 5-1), the generation of surrogate virtual cohorts of arbitrarily large size could provide natural candidates for future data augmentation applications.

Yet, by capitalizing exclusively on redundancy, augmentation cannot replace the gathering of more empirical data (Carrillo et al., 2012; Toga et al., 2016). Unfortunately, federation (or even mining) of data is often impeded by unavoidable juridical concerns linked to strict and diverse regulations (Dulong de Rosnay, 2017; Thorogood et al., 2018) The use of virtual cohorts may once again relieve this burden. Virtual cohorts maintain their statistical relation to the original data, in a way sufficiently good to be exploitable for classification, but do not precisely match the original data, maintaining an inherent variability. This fact may constitute a feature rather than a bug, in the context of data sharing. Indeed, if virtual data carry information operationally equivalent to the one carried by empirical data, they do not carry exactly the same information. It is not, therefore, possible to exactly reconstruct the original subject data from virtualized connectomes, and privacy concerns would be considerably reduced if not entirely removed by sharing dual space images of actual data –eventually demultiplied into virtual cohorts– rather than the original real space data. We thus anticipate a near future in which virtual cohorts, providing vast numbers of virtual and bi-virtual connectivity information, will play an increasing role in massive data-driven explorations of factors predictive of pathology and, in particular, neurodegenerative disease progression.

## Acknowledgements

Data collection and sharing for this project was funded by the Alzheimer’s Disease Neuroimaging Initiative (ADNI) (National Institutes of Health Grant U01 AG024904) and DOD ADNI (Department of Defense award number W81XWH-12-2-0012). ADNI is funded by the National Institute on Aging, the National Institute of Biomedical Imaging and Bioengineering, and through generous contributions from the following: AbbVie, Alzheimer’s Association; Alzheimer’s Drug Discovery Foundation; Araclon Biotech; BioClinica, Inc.; Biogen; Bristol-Myers Squibb Company; CereSpir, Inc.; Cogstate; Eisai Inc.; Elan Pharmaceuticals, Inc.; Eli Lilly and Company; EuroImmun; F. Hoffmann-La Roche Ltd and its affiliated company Genentech, Inc.; Fujirebio; GE Healthcare; IXICO Ltd.; Janssen Alzheimer Immunotherapy Research & Development, LLC.; Johnson & Johnson Pharmaceutical Research & Development LLC.; Lumosity; Lundbeck; Merck & Co., Inc.; Meso Scale Diagnostics, LLC.; NeuroRx Research; Neurotrack Technologies; Novartis Pharmaceuticals Corporation; Pfizer Inc.; Piramal Imaging; Servier; Takeda Pharmaceutical Company; and Transition Therapeutics. The Canadian Institutes of Health Research is providing funds to support ADNI clinical sites in Canada. Private sector contributions are facilitated by the Foundation for the National Institutes of Health (www.fnih.org). The grantee organization is the Northern California Institute for Research and Education, and the study is coordinated by the Alzheimer’s Therapeutic Research Institute at the University of Southern California. ADNI data are disseminated by the Laboratory for Neuro Imaging at the University of Southern California. DB acknowledges support from the EU Innovative Training Network “i-CONN” (H2020 ITN 859937) and VJ acknowledges funding by the European Union’s Horizon 2020 Framework Program for Research and Innovation under the Specific Grant Agreement No. 785907 (Human Brain Project SGA2) and H2020 Research and Innovation Action grants VirtualBrainCloud, and RM acknowledges Brightfocus Foundation ADR grant program, grant reference number: A2017286S.

## Extended data

Extended Data 1. MATLAB® workspaces including virtual SC and FC connectomes generated with our data completion pipelines as well as virtual cohorts. All workspaces are available at the address https://github.com/FunDyn/VirtualCohorts.

**Extended Data Table 1-1.**
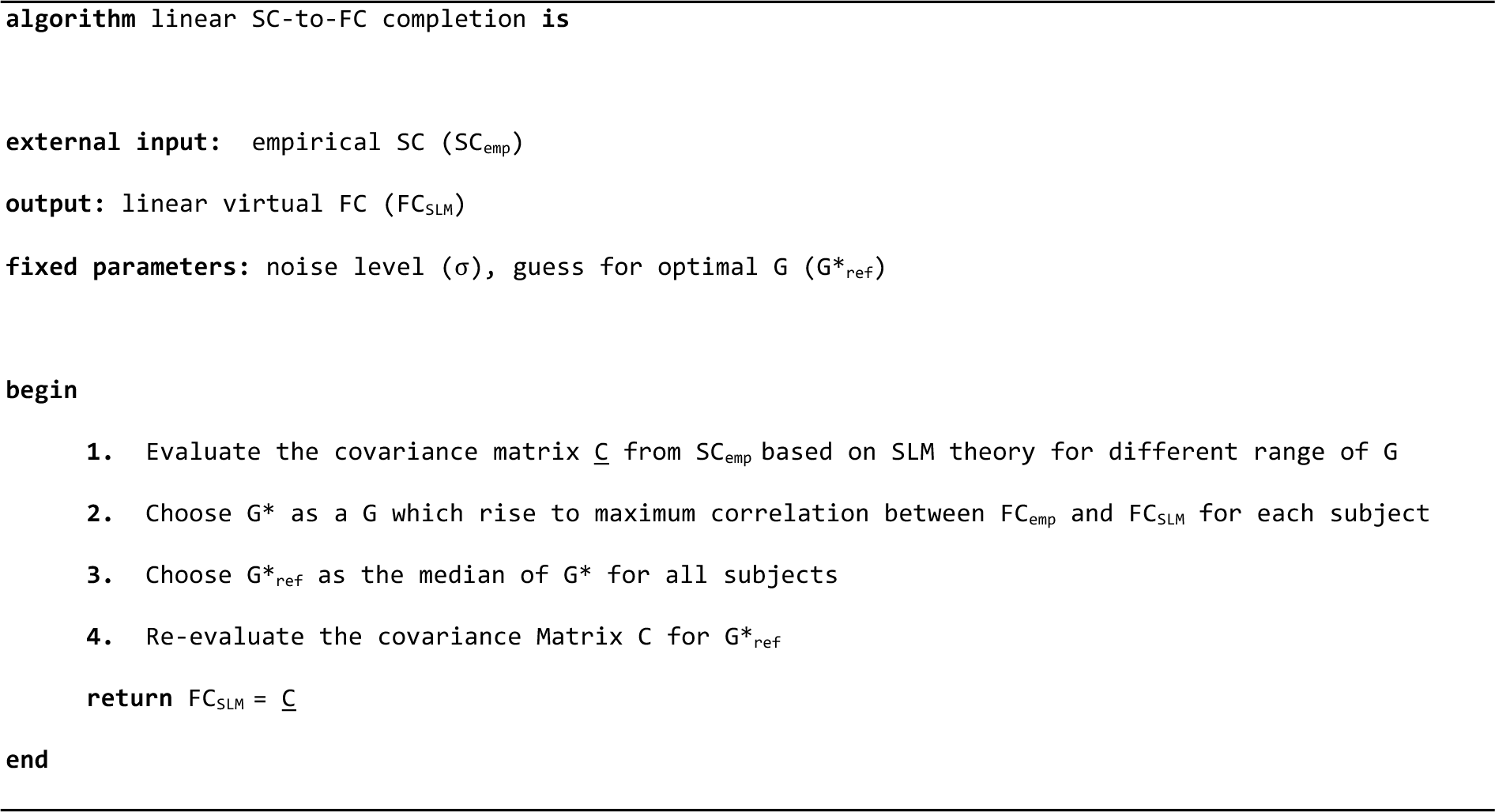
Pseudo-code for linear SC-to-FC completion

**Extended Data Table 2-1.**
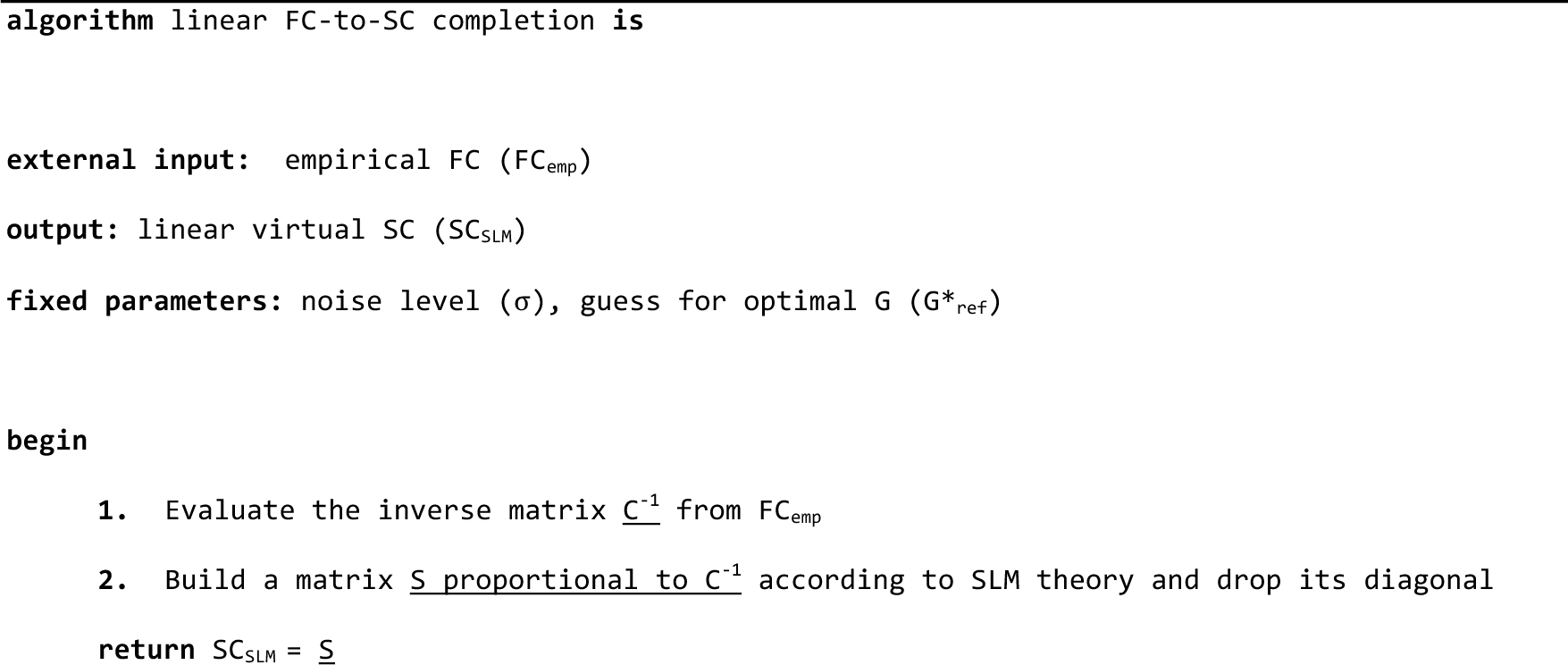
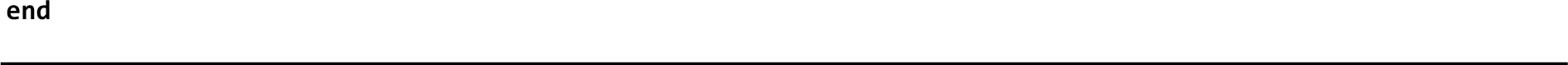
Pseudo-code for linear FC-to-SC completion

**Extended Data Table 3-1.** Discriminating control and patient subjects in the ADNI subset with only SC connectomes.

| *Tested on*<br>*Trained on* | SC<sub>emp</sub> | FC<sub>MFM</sub> | SC<sub>biMFM</sub> |
| --- | --- | --- | --- |
| SC<sub>emp</sub> | 0.69 [0.61 < AUC < 0.74] | n.s. | 0.55 [0.51 < AUC < 0.60] |
| FC<sub>MFM</sub> | 0.54 [0.50 < AUC < 0.59] | 0.64 [0.57 < AUC < 0.69] | 0.62 [0.54 < AUC < 0.68] |
| SC<sub>biMFM</sub> | 0.56 [0.51 < AUC < 0.62] | 0.62 [0.53 < AUC < 0.68] | 0.59 [0.52 < AUC < 0.64] |

*Indicated values are median and 5% and 95% percentiles over crossvalidation replicas of the indicated classification.*

**Extended Data Table 3-2.** Discriminating control and patient subjects in the ADNI subset with only FC connectomes.

| *Tested on*<br>*Trained on* | FC<sub>emp</sub> | SC<sub>MFM</sub> | FC<sub>biMFM</sub> |
| --- | --- | --- | --- |
| FC<sub>emp</sub> | 0.75 [0.70 < AUC < 0.79] | 0.71 [0.65 < AUC < 0.79] | 0.65 [0.58 < AUC < 0.71] |
| SC<sub>MFM</sub> | 0.69 [0.61 < AUC < 0.75] | 0.73 [0.67 < AUC < 0.78] | 0.55 [0.50 < AUC < 0.61] |
| FC<sub>biMFM</sub> | 0.70 [0.65 < AUC < 0.76] | n.s. | n.s. |

*Indicated values are median and 5% and 95% percentiles over crossvalidation replicas of the indicated classification.*

**Extended Data Table 5-1.** Inter-subject distances for empirical – bivirtual pairs.

| Type of completion |  | Inter-distance correlation |  |
| --- | --- | --- | --- |
|  |  | ADNI | Healthy aging |
| SCemp to SCemp vs | SCbiSLM to SCbiSLM | 0.39*** [0.37, 0.41] | 0.81*** [0.79, 0.83] |
|  | SCbiMFM to SCbiMFM | 0.39*** [0.36, 0.43] | 0.53*** [0.48, 0.57] |
| FCemp to FCemp vs | FCbiSLM to FCbiSLM | 0.06*** [0.5, 0.8] | 0.55*** [0.50, 0.58] |
|  | FCbiMFM to FCbiMFM | 0.43*** [0.42, 0.44] | 0.40*** [0.36, 0.44] |

*Indicated values are median and 5% and 95% percentiles over bootstrap with replacement replicas of correlation computation*

**Extended Data Figure 2-1.**
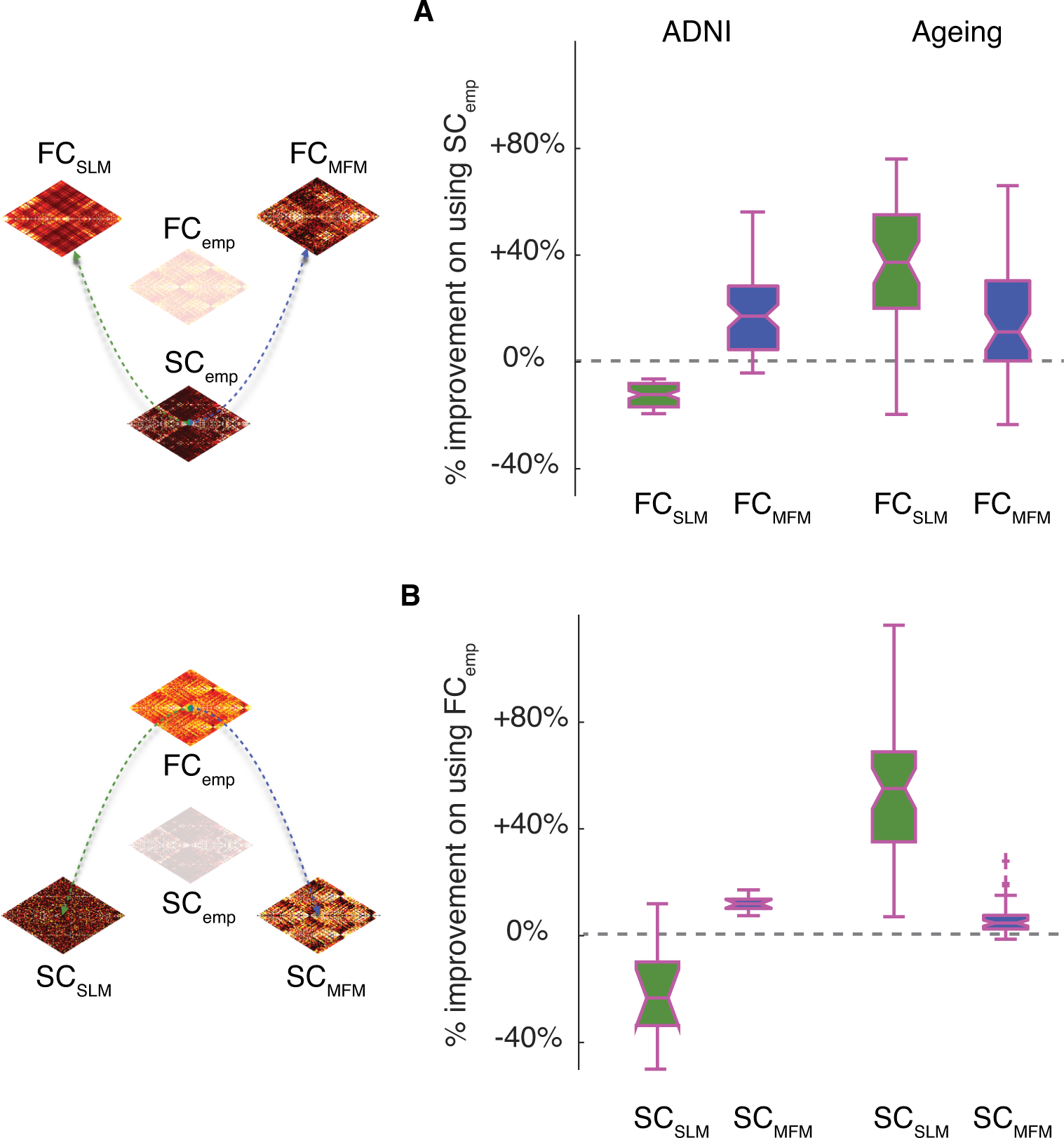
Viability of data completion. We checked whether the performance of data completion based on the algorithmic procedures of Tables 1 and 2 or 1-1 and 2-1 is superior to the one of a trivial strategy in which the target connectome to reconstruct is just taken to be identical to the “other connectome” (i.e. using SC, when trying to reconstruct missing FC; or using FC, when trying to reconstruct missing SC). A-B) We computed percent improvement in data completion over the trivial “other connectome” strategy using a SLM-based or an MFM-based data completion method, focusing on the “SC_emp_ + FC_emp_” subset for which both ground truth connectomes are known. A) Percent improvements in data completion when completing FC from SC. B) Percent improvements in data completion when completing SC from FC. For the SLM-based functional data completion approach, the use of FC_SLM_ on the ADNI dataset resulted in a worse performance (median drop Δ_trivial_ = -15%, see Materials and Methods for definition), however, for the healthy ageing dataset the use of FC_SLM_ resulted in a clearly better performance than when using “the other connectome” (median improvement Δ_trivial_ = +40%); similarly, applying the SLM-based approach for the structural data completion, the use of SC_SLM_ rather than FC_emp_ as an ersatz for SC_emp_ leads to drops of improvements in quality with a median value of approximately -20%, for the ADNI dataset but an increase of nearly ∼50% for the healthy aging dataset. Thus, the performance of linear data completion can yield to good results, but this performance did not robustly generalize through datasets. On the other hand, for the MFM-based functional data completion, the median improvement was close to ∼20% for both datasets which can go as high as +60% in some subjects; using the same approach but for the structural data completion, the performance was lower than non-linear SC-to-FC data completion, with median improvement of ∼15% for the ADNI dataset and of ∼10% for the healthy aging dataset.

**Extended Data Figure 3-1.**
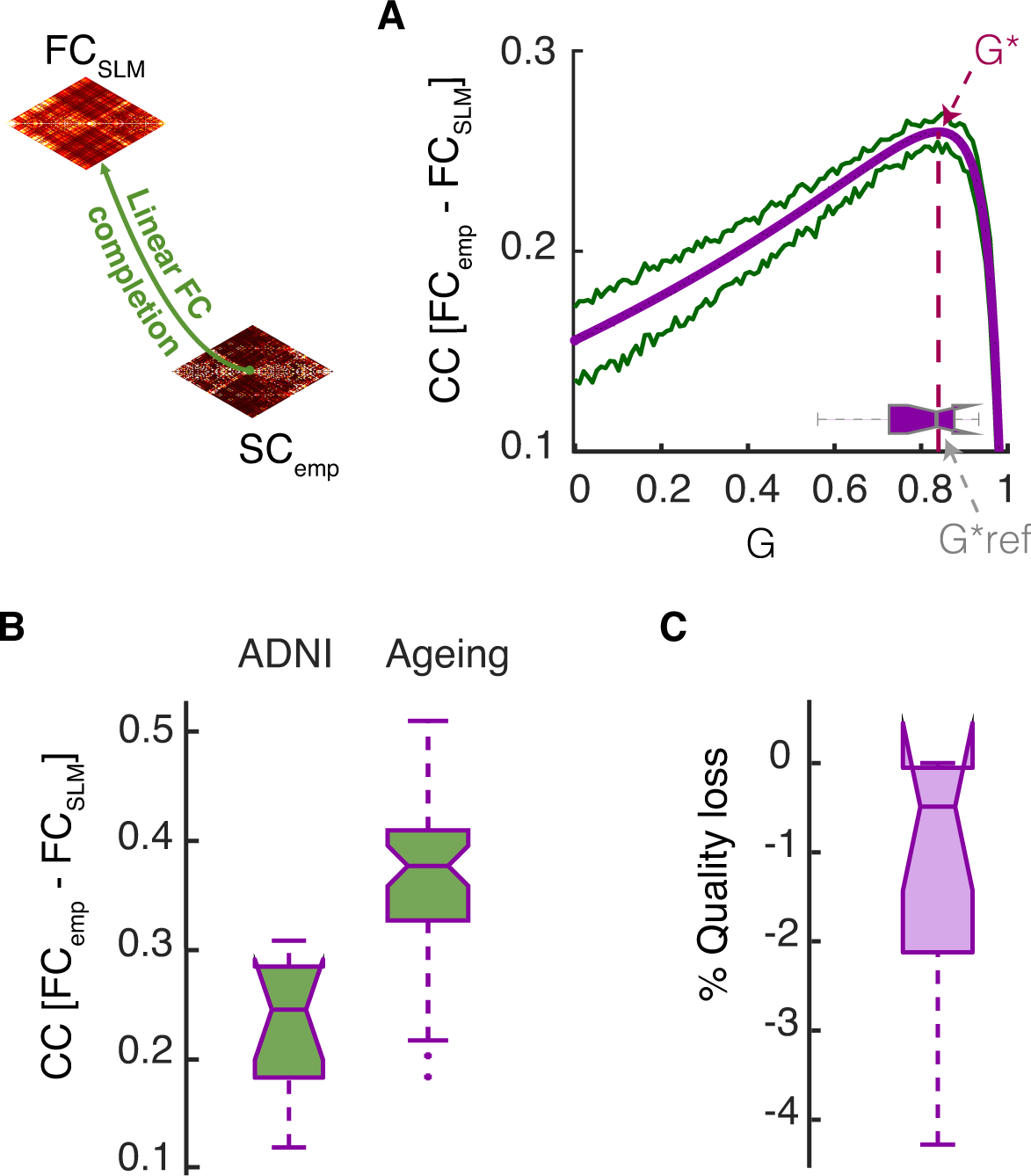
Linear SC-to-FC data completion. The functional data completion can also be done using the linear model starting from SC_emp_ matrices. A) the systematic exploration (for a representative subject) of the dependency of correlation between FC_emp_ and FC_SLM_ on the SLM parameter G (global scale of long-range connectivity strength) shown by the violet line indicates that the best fitting value *G\** (dashed line) can be obtained slightly before the critical point of the system *G_critic_* = 1/*max*(λ_*i*_) which since the SC_emp_ matrices are normalized to one 1/*max*(λ_*i*_) = 1 and *G_critic_* = 1. The green lines display 5 and 95 percentiles of bootstrap resampling. The inset boxplot gives the distribution of *G\** over all the subjects in the “SC_emp_ + FC_emp_” subset; for the SLM SC-to-FC completion, we used a common value *G*_ref_* = 0.83, equal to the median of the boxplot. B) The boxplot reports the distribution of Pearson correlation between FC_emp_ and FC_SLM_ for all subjects from the “SC_emp_ + FC_emp_” subset with a median equal to 0.243 for the ADNI dataset and 0.377 for the Healthy Ageing dataset. C) In case of using the common value *G*_ref_* for all subjects instead of the actual personalized optimum *G\** for each subject in the “SC_emp_ + FC_emp_” subset, the value of quality loss for each subject is shown in the boxplot with median equal to 0.5%.

**Extended Data Figure 3-2.**
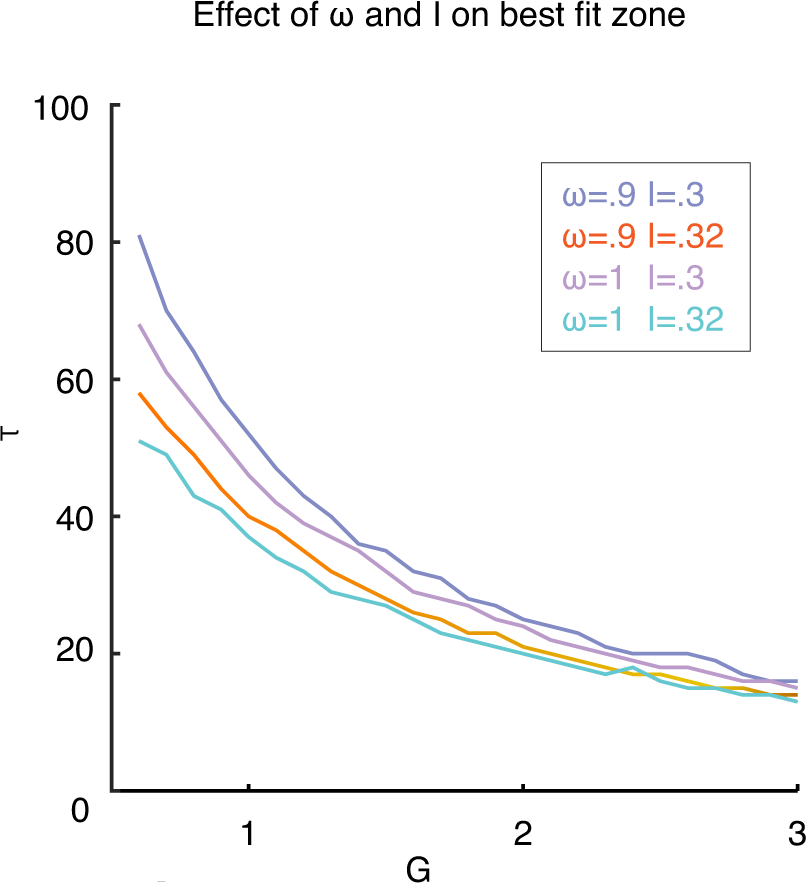
The dependency of best MFM fit zone on additional regional dynamics parameters. In the non-linear data completion, the global parameters of the MFM model are *G* (inter-regional coupling strength), *τ* (synaptic time-constant of within-region excitation), *ω* (relative strength of recurrent within-region connections) and *I* (external input) which parameters *G* and *τ* were investigated in this paper (see Figure 3). Here we showed for different values of *ω* and *I*, the narrow concave stripe of Figure 3.A as the representative of the best fitting zone is slightly shifted in the G/*τ* plane, suggesting G and *τ* are more sensitive parameters and need to be explored rather than *ω* and *I*.

**Extended Data Figure 4-1.**
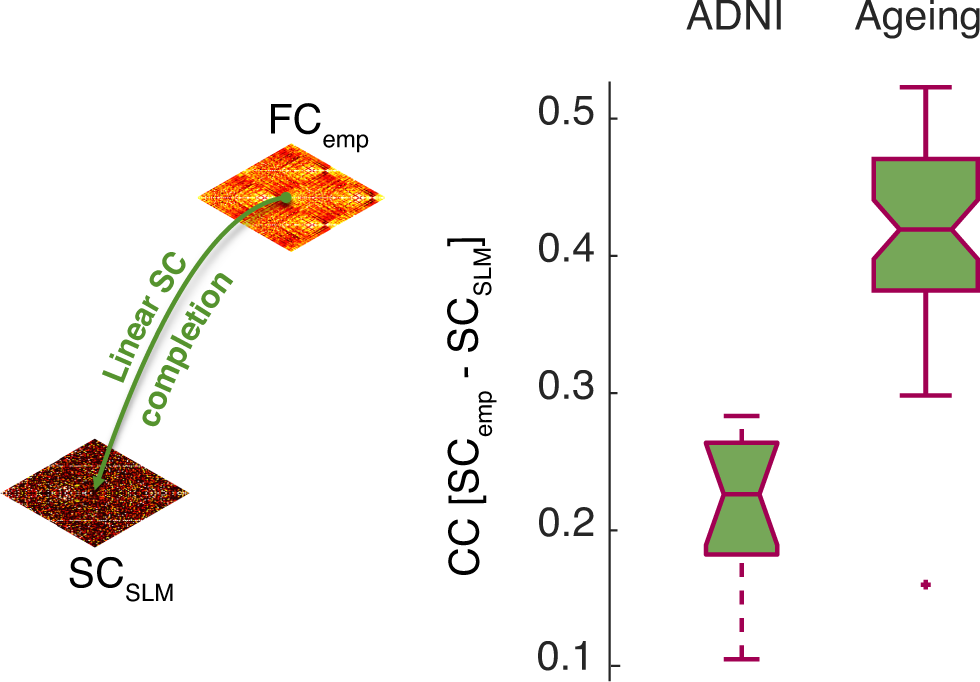
Linear FC-to-SC data completion. Using the linear model, it is equivalently possible to infer the structural SC_SLM_ matrices from FC_emp_. Since in this approach the free parameters of SLM model appear as scaling factor, they don’t affect the correlation of the inferred SC_SLM_ with the SC_emp_ so there is no need for parameter exploration here. The distribution of the correlation values for all the subjects from the “SC_emp_ + FC_emp_” ADNI subset is shown in the boxplot with median equal to 0.21 and 0.42 for the Healthy Ageing dataset.

**Extended Data Figure 6-1.**
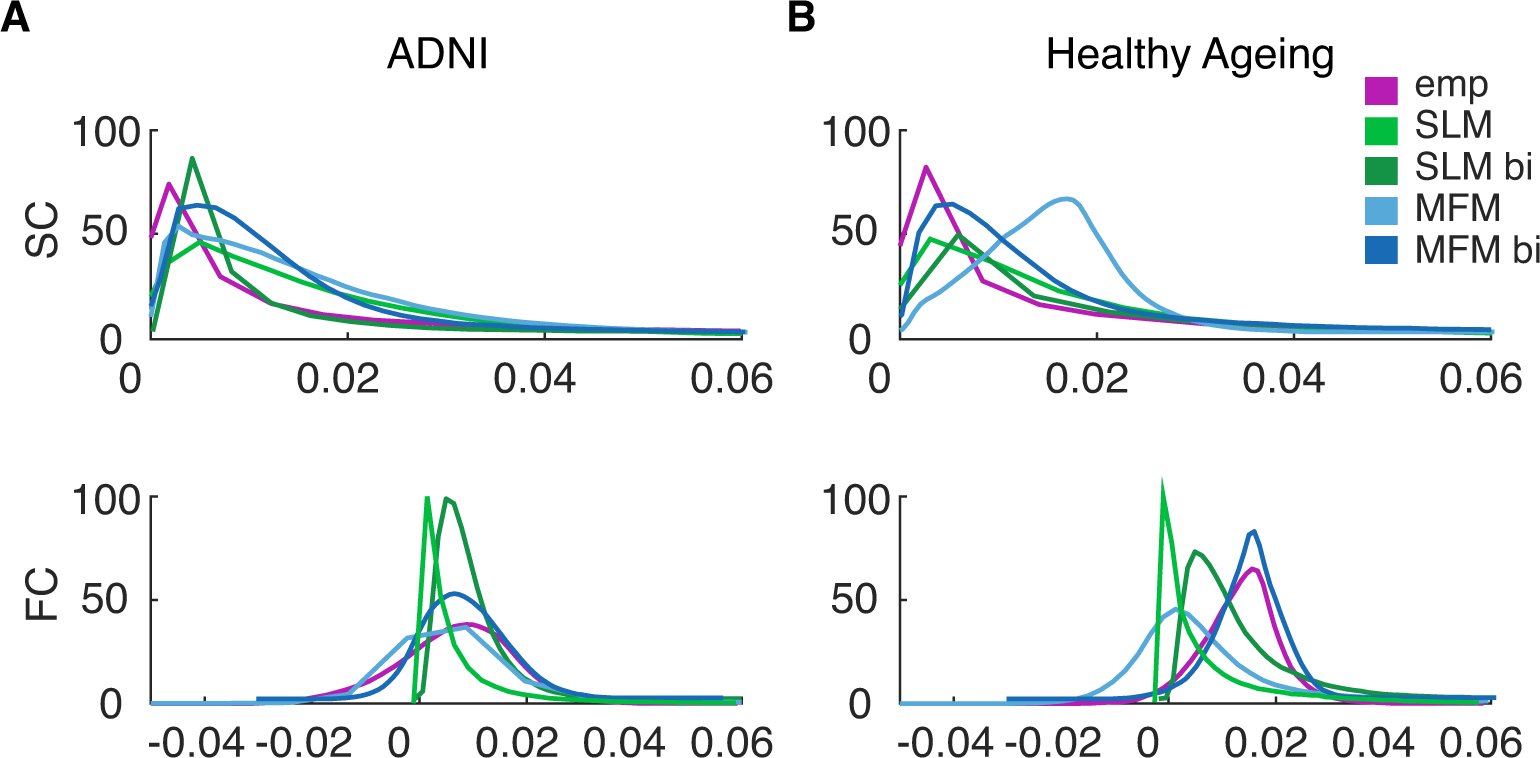
The global distributions of link weights for all different types of connectomes. Most of the distributions show similarity in shape with their empirical counterparts (pink). SC weights distributions with a peak for small values and a fat right tail; FC weights distributions with more symmetric and a broader peak at intermediate strengths.

**Extended Data Figure 7-1.**
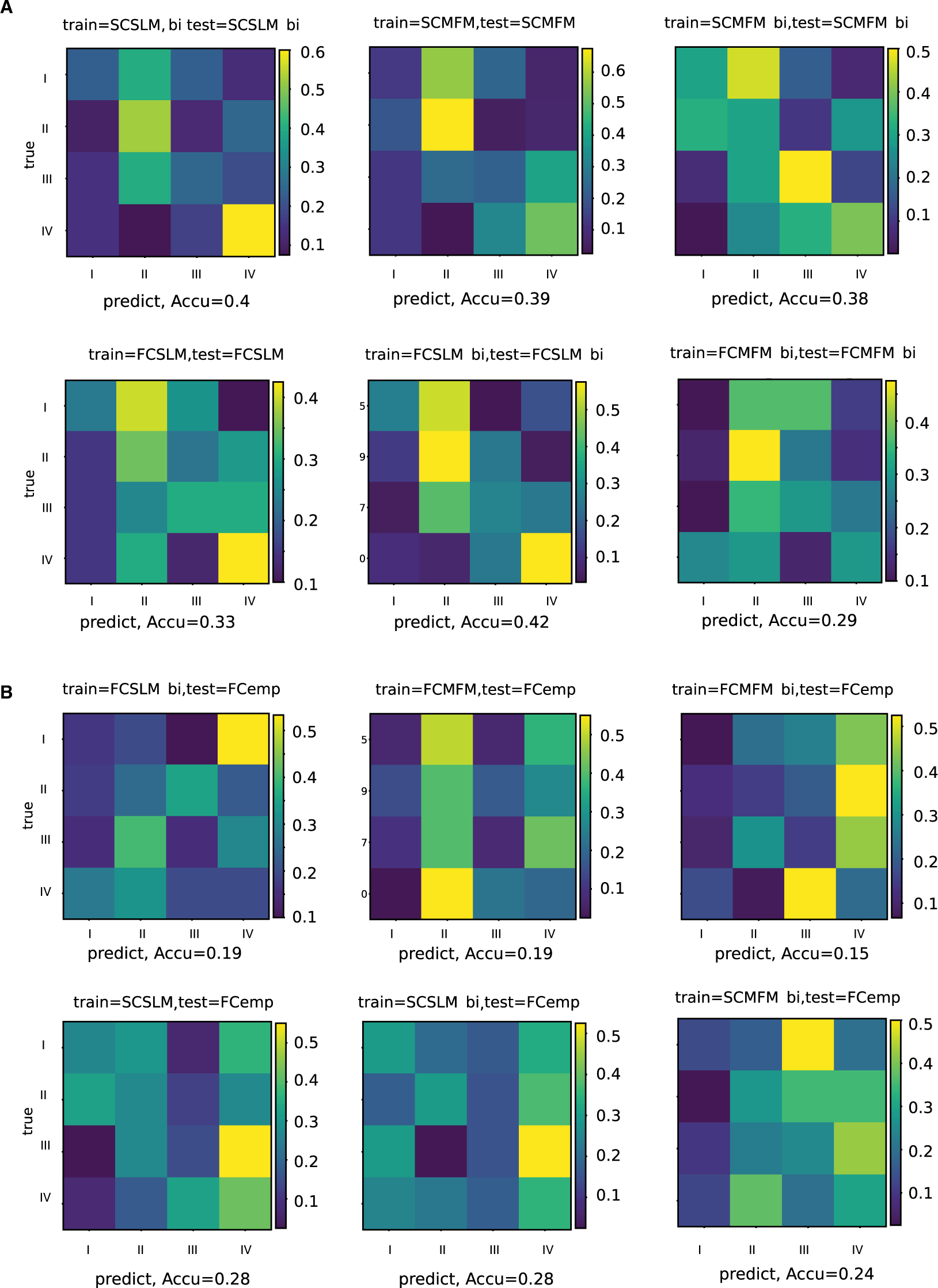
Age class discriminations on the healthy ageing dataset. A) The classification performances when the classifier was train and tested on the same virtual connectome is above chance level (∼0.25) with maximum accuracy of ∼0.42 for FC_SLM-bi_ and minimum accuracy of ∼0.29 for FC_MFM-bi_. B) The classification accuracy dropped when the classifier was trained on the virtual connectome and tested on the empirical connectome. The only cases where the accuracy was above chance level was when the classifier was trained on SC_SLM_ and SC_SLM-bi_ and tested on FC_emp_ connectome, with an accuracy of ∼0.28.

**Extended Data Figure 8-1.**
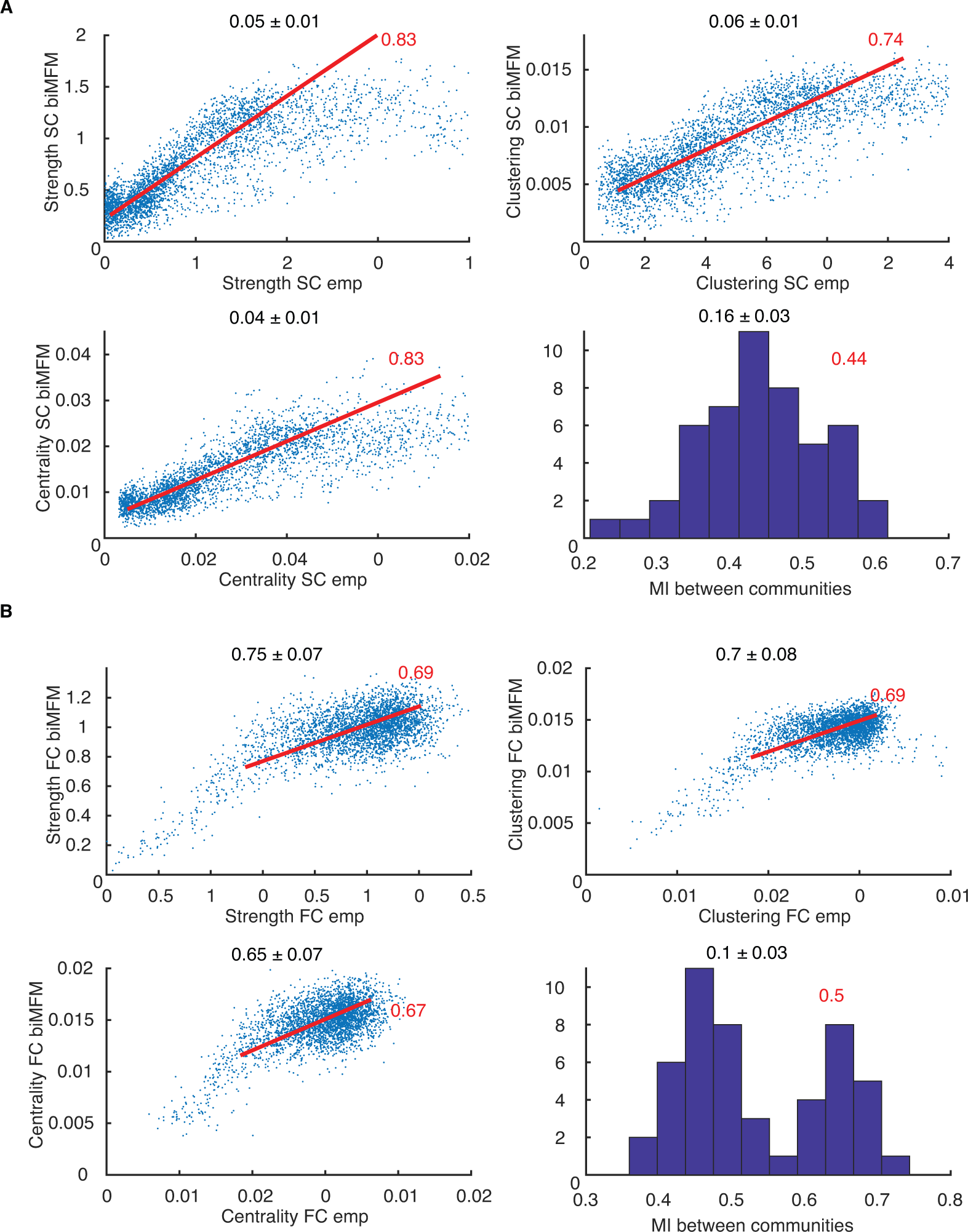
Correspondence of network topology between empirical and their bivirtual dual connectomes (healthy aging dataset). We show here scatter plots of connectivity strengths (top left), local clustering coefficients (top right) and local centrality coefficients (bottom left) for different brain regions and subjects, plotting feature values for empirical connectomes vs their bivirtual counterparts and the histograms over different subjects of the relative mutual information (normalized between 0 and 1, the latter corresponding to perfect matching) between the community structures (bottom right) of empirical connectomes and their bivirtual duals. Results are shown in panel A for SC and in panel B for FC connectomes for the healthy ageing dataset (see Figure 8 for the comparison with the ADNI dataset). Again for both cases, we see a remarkable correlation at the ensemble level between network topology features for empirical bivirtual connectomes (see Table 4 for the superior correspondence at the single subject level for the ageing dataset).

